# Precise, Fast and Comprehensive Analysis of Intact Glycopeptides and Modified Saccharide Units with pGlyco3

**DOI:** 10.1101/2021.02.06.430063

**Authors:** Wen-Feng Zeng, Wei-Qian Cao, Ming-Qi Liu, Si-Min He, Peng-Yuan Yang

## Abstract

We present a glycan-first glycopeptide search engine, pGlyco3, to comprehensively analyze intact N- and O-glycopeptides, including glycopeptides with modified saccharide units. A novel glycan ion-indexing algorithm developed in this work for glycan-first search makes pGlyco3 5-40 times faster than other glycoproteomic search engines without decreasing the accuracies and sensitivities. By combining electron-based dissociation spectra, pGlyco3 integrates a fast, dynamic programming-based algorithm termed pGlycoSite for site-specific glycan localization (SSGL). Our evaluation based on synthetic and natural glycopeptides showed that the SSGL probabilities estimated by pGlycoSite were proved to be appropriate to localize site-specific glycans. With pGlyco3, we found that N-glycopeptides and O-mannose glycopeptides in yeast samples were extensively modified by ammonia adducts on Hex (aH) and verified the aH-glycopeptide identifications based on released N-glycans and ^15^N/^13^C-labeled data. Thus pGlyco3, which is freely available on https://github.com/pFindStudio/pGlyco3/releases, is an accurate and flexible tool to identify glycopeptides and modified saccharide units.

## Introduction

Protein glycosylation is a fundamental posttranslational modification (PTM) that is involved in many biological functions^1-3^. In recent years, tandem mass spectrometry (MS/MS) has been shown to be a promising technique to analyze site-specific glycans on proteins^4, 5^. Modern MS instruments have integrated different fragmentation techniques for glycopeptide analysis, such as higher-energy collisional dissociation (HCD), electron-transfer dissociation (ETD), electron-transfer/higher-energy collision dissociation (EThcD), and electron-transfer/collision-induced dissociation (ETciD)^6, 7^. HCD, especially stepped collision energy HCD (sceHCD), can provide abundant glycopeptide Y ions (glycan Y ions with an intact peptide attached) and quite a few b/y ions of naked peptides to identify the glycan parts and peptide parts, respectively^8, 9^. Information of Y ions allows us to directly identify the entire glycan composition, and core Y ions (e.g., Y0, Y1, and Y2) can be used to determine the glycan and peptide masses. The b/y ions and b/y+HexNAc ions in HCD can be used to identify peptide parts, but they commonly do not provide enough information to determine site-specific glycans for multiple glycosylated sites with a given sceHCD spectrum. ETxxD (ETD, EThcD, and ETciD) could generate glycan-attached c/z ions to not only identify peptides but also deduce site-specific glycans^10-13^.

Based on modern MS techniques, many glycopeptide search engines have been developed over the last decade^4, 5, 14^. There are three main search strategies for intact glycopeptide identification: peptide-first, glycan-removal, and glycan-first. The peptide-first search is arguably the most widely used strategy and was adopted by Byonic^15^, gpFinder^16^, GPQuest^17^, pMatchGlyco^18^, GPSeeker^19^, Protein Prospector^11^, and two recently developed tools MSFragger (MSFragger-Glyco^20^) and MetaMorpheus (MetaMorpheus O-Pair^21^). This method first searches the peptide part and then deduces the glycan part as a large variable modification by considering some B/Y ions. MSFragger and MetaMorpheus use the peptide ion-indexing technique^22^ to accelerate the peptide-first search. The glycan-removal search deduces the pseudopeptide masses from N-glycopeptide spectra by using potential reducing-end Y ions, and then it modifies the spectral precursor masses as the pseudopeptide masses to identify peptides using conventional peptide search engines^23-25^. Based on the deduced glycan mass, MAGIC searches the glycan compositions by using the knapsack algorithm without glycan composition/structure databases^25^. The recently developed O-search further extended the glycan-removal strategy for O-glycopeptide identification^26^. These tools, especially Byonic, MSFragger and MetaMorpheus, have increased the identification sensitivity for glycoproteomics^27^; however, glycan-level quality control is not a priority. Glycan-first search is used in the pGlyco software series^8, 28^ as well as other tools (Sweet-Heart^29^ and GlycoMaster DB^30^), and it first searches the glycan parts to remove unreliable glycans and then searches the peptide parts. pGlyco 2.0 is the first search engine that can perform glycan-, peptide- and glycopeptide-level quality control for glycopeptides. It was extended for peptide identification and tandem mass tag (TMT)-quantification with MS3 by SugarQuant^31^. However, pGlyco 2.0 supports only the search for normal N-glycans of mammals in GlycomeDB^32^ with sceHCD spectra, and hence, it is difficult for users to apply to analyze customized glycans or modified saccharide units (e.g., phospho-Hex or mannose-6-phosphate). Furthermore, modern glycopeptide search engines should consider site-specific glycan localization (SSGL), as ETxxD techniques have been widely used in glycoproteomics. Some works, such as the “Delta Mod score” of Byonic and “SLIP”^33^ of Protein Prospector, extended the site localization algorithms from traditional PTMs to glycosylation, and assessed the localization reliabilities. However, traditional PTM search algorithms do not address the computational complexity of SSGL for glycopeptides, which has been well discussed by Lu et al.^21^ Graph-based SSGL algorithms, such as GlycoMID for hydroxylysine O-glycosylation^34^ and MetaMorpheus for common O-glycosylation^21^, enabling the fast localization process. However, accurate SSGL and its validation are still unresolved problems.

Here, we proposed pGlyco3, a novel glycopeptide search engine that enables analysis of modified saccharide units and SSGL. 1) pGlyco3 applies the glycan-first search strategy to accurately identify glycopeptides. It uses canonicalization-based glycan databases to support the modified saccharide unit analysis and a novel glycan ion-indexing technique to accelerate the search for glycans. 2) For SSGL, we developed a dynamic programming algorithm termed pGlycoSite to efficiently localize the site-specific glycans using ETxxD spectra. 3) We emphasized validation for glycopeptide identification and SSGL. To validate the accuracy of pGlyco3, we designed several experiments to show that pGlyco3 outperformed other tools in terms of the identification accuracies for both N- and O-glycopeptides, especially at the glycan level. We also designed four methods to validate the SSGL of pGlycoSite. Our validations showed that SSGL probabilities estimated by pGlycoSite were proved to be appropriate to localize site-specific glycans. 4) We used pGlyco3 to identify a modification on Hex (Hex with an ammonia adduct, simplified as “aH”) on N-glycopeptides and O-mannose (O-Man) glycopeptides in yeast samples. We validated the aH search results on yeast with N-glycome data and ^15^N/^13^C-labeled glycopeptide data. This analysis further demonstrated the reliability and flexibility of pGlyco3 for intact glycopeptide and modified saccharide unit identification.

## Results

### Workflow of pGlyco3

pGlyco3 uses sceHCD and ETxxD spectra to identify glycopeptides, analyze modified saccharide units, estimate glycan/peptide false discovery rates (FDRs), and localize site-specific glycans (Fig. 1a). For HCD-pd-ETxxD spectra, pGlyco3 merges HCD and ETxxD spectra before searching. The whole workflow is described in the Online Methods. For glycan part identification, pGlyco3 provides several built-in N- and O-glycan databases, and it can generate new glycan databases from GlycoWorkbench^35^ with expert knowledge. Benefitting from the flexible canonicalization-based glycan representation, pGlyco3 enables the convenient analysis of modified or labeled glycans of glycopeptides (Online Methods, Supplementary Note 1).

**Figure 1.**
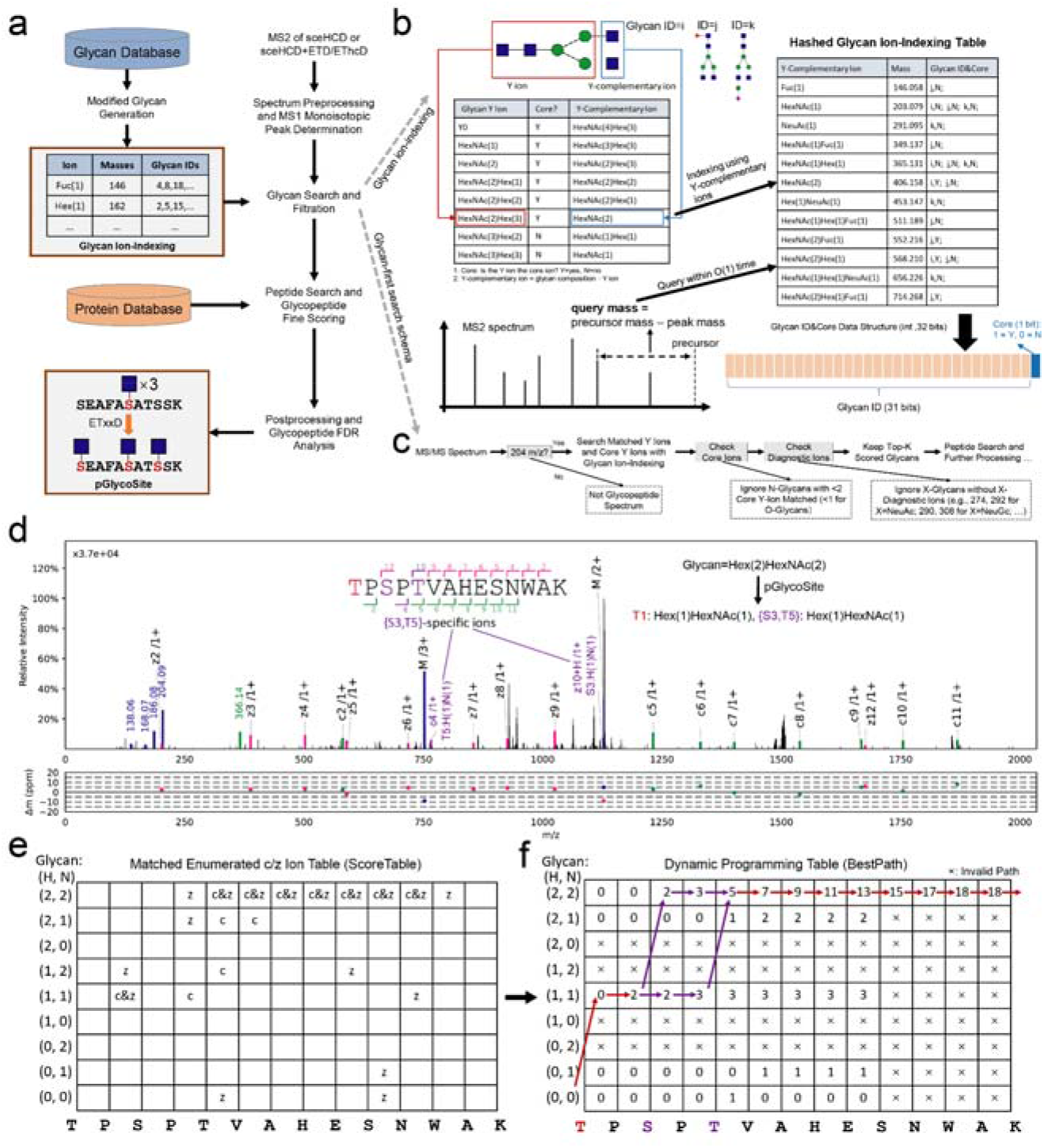
Workflow of pGlyco3 and its algorithms. (a) Software schema. (b) Illustration of the glycan ion-indexing technique. The ion-indexing information of glycan j and k is displayed in Supplementary Note Fig. 3. (c) Glycan ion-indexing-based glycan-first search schema of pGlyco3. (d-f) Representative example of pGlycoSite. (d) An EThcD-GPSM (“TPSPTVAHESNWAK + H(2)N(2)”, H=Hex, N=HexNAc) with localized sites using the pGlycoSite algorithm. (e) shows all possible matched c/z ions against the EThcD spectrum (*ScoreTable*). (f) shows the dynamic programming table from bottom left to top right (*BestPath*). The arrow lines indicate the best-scored paths, and the purple arrow lines show that two paths share the same score from S3 to T5. T1 is uniquely localized with Hex(1)HexNAc(1), and {S3:T5} is localized as a “site-group” with Hex(1)HexNAc(1), as shown in (d). Details on calculation of the *ScoreTable* and *BestPath* are illustrated in the Online Methods and Supplementary Note 3.

In contrast to Byonic, MetaMorpheus and MSFragger, pGlyco3 applies the glycan-first strategy as it first searches the glycan parts and filters out unreliable ones. For a fast glycan-first search, we designed a glycan ion-indexing technique to score all glycans as well as their core Y ions (core Y ions are defined in Supplementary Table 2) by matching the query spectrum only once within the linear search time (i.e., O(#peaks), Supplementary Note 2). The key observation of glycan ion-indexing is that if a peak is a Y ion of a glycopeptide, then the precursor mass minus the peak mass would be the Y-complementary ion mass (Online Methods). As the Y-complementary ion mass does not contain the peptide mass, it enables us to search the glycans before peptides are identified. Therefore, the ion-indexing of glycans in pGlyco3 indexes Y-complementary masses instead of Y ions, enabling the fast glycan-first search (Fig. 1b, Online Methods, Supplementary Note 2). As shown in Fig. 1b, the hashed keys of the glycan ion-indexing table are the Y-complementary ion masses, and the values are the list of glycan IDs from which the Y-complementary ions originate. As the core Y ions are very important in glycan identification, we use an extra bit in the glycan ID of the indexing table to indicate whether the corresponding Y ion of the Y-complementary ion is a core ion. Note that the Y0 ion is always considered as a core ion for an N- or O-glycopeptide in pGlyco3. The schema of the glycan-first search is shown in Fig. 1c.

This method applies a few verification steps to ensure the reliability of the remaining glycans, including the number of matched core Y ions, the existence of glyco-specific diagnostic ions, and the rank of glycan scores. After glycan filtration, pGlyco3 performs the peptide search, glycan/peptide fine-scoring, postsearch processing, and glycopeptide FDR estimation (Fig. 1a).

pGlyco3 integrates a novel algorithm termed pGlycoSite to localize the site-specific glycans and estimate the localization probabilities using c/z ions in ETxxD spectra. For a given spectrum with the identified peptide and the glycan composition, the complexity of enumerating all possible glycopeptide forms will exponentially increase as the number of candidate sites and monosaccharides increase (Online Methods, Supplementary Note 3). Instead of generating the glycopeptide forms, pGlycoSite enumerates a table of all possible c/z ions, matches the table against the ETxxD spectrum. Although the number of glycopeptide forms may be exponentially large, there are only *F* × (*L* − 1) possible c or z ions, where *F* is the number of subglycan compositions and *L* is the peptide length. The pGlycoSite algorithm is extremely fast, as its computational complexity is *O* (*L*, × *F*^2^) (Online Methods). Fig. 1d-f show an exmaple of how pGlycoSite works. Based on the matched c/z ion table (*ScoreTable* in Fig. 1e), pGlycoSite uses a dynamic programming algorithm to obtain the best-scored path across the table from bottom left to top right. If multiple paths reach the same best score, pGlycoSite will regard the amino acids from the branching position to the merging position as a “site-group”. As shown in Fig. 3f, T1 with Hex(1)HexNAc(1) is uniquely localized, and {S3:T5} is a “site-group” because either S3 or T5 has a supporting site-specific c/z ion (Fig. 3d), resulting in the same score. More details of the pGlycoSite algorithm, including calculation of the *ScoreTable, BestPath* and localization probability estimation, are illustrated in the Online Methods and Supplementary Note 3. After the SSGL probabilities are estimated, the SSGL-FDR can be deduced and used to validate the accuracies of the estimated SSGL probabilities (Online Methods).

### pGlyco3 for N-glycopeptide identification

To demonstrate the performance of pGlyco3, we comprehensively compared pGlyco3 with Byonic, MetaMorpheus, and MSFragger using two N-glycopeptide datasets.

To compare the precision of identified glycopeptides, we used our previously published data of unlabeled, ^15^N-labeled, and ^13^C-labeled fission yeast mixture samples (PXD005565^8^) to test these software tools. The searched protein database was concatenated proteome database of fission yeast and mouse, and the searched N-glycan database contained NeuAc-glycans which should not be identified in yeast samples. The search details are listed in the Online Methods and Supplementary Data 1. We analyzed three levels of identification errors: 1) The element-level error. The unlabeled glycopeptide-spectrum matches (GPSMs) may be identified with incorrect numbers of N or C elements if the GPSMs could not be verified by ^15^N- or ^13^C-labeled MS1 precursors; 2) The glycan-level error. The glycan parts tend to be incorrectly identified if the GPSMs contain NeuAc-glycans; 3) The peptide-level error. The peptide parts are false positives if they are from the mouse protein database. The testing results are shown in Fig. 2a. All tools showed low peptide-level error rates, implying that both peptide- and glycan-first searches are accurate at identifying the peptide parts of glycopeptides. However, the peptide-first-based tools showed high glycan-level error rates for yeast glycopeptides. MSFragger yielded the most identified GPSMs, but 24.1% contained NeuAc-glycans. The percentages of NeuAc-glycans obtained by MetaMorpheus and Byonic were 7.0% and 15.3%, respectively. On the other hand, benefitting from the essential glycan ion analyses and glycan FDR estimation, pGlyco3 showed good performance in controlling element-, glycan-, and peptide-level error rates without losing the number of identified GPSMs. This does not mean that pGlyco3 is 100% accurate at the glycan level even if there are no NeuAc-identifications. With different glycan databases, pGlyco3’s glycan-level error rates would be 0.8-4% (Supplementary Note 5), but it was still much better than the others. This comparison does not imply that the peptide-first strategy is not accurate. Instead, it suggests that glycan-level quality control is also necessary for the peptide-first search. pGlyco3 also showed good performance for glycopeptide identification on HCD-pd-EThcD spectra, as shown in Supplementary Fig. 7.

**Figure 2.**
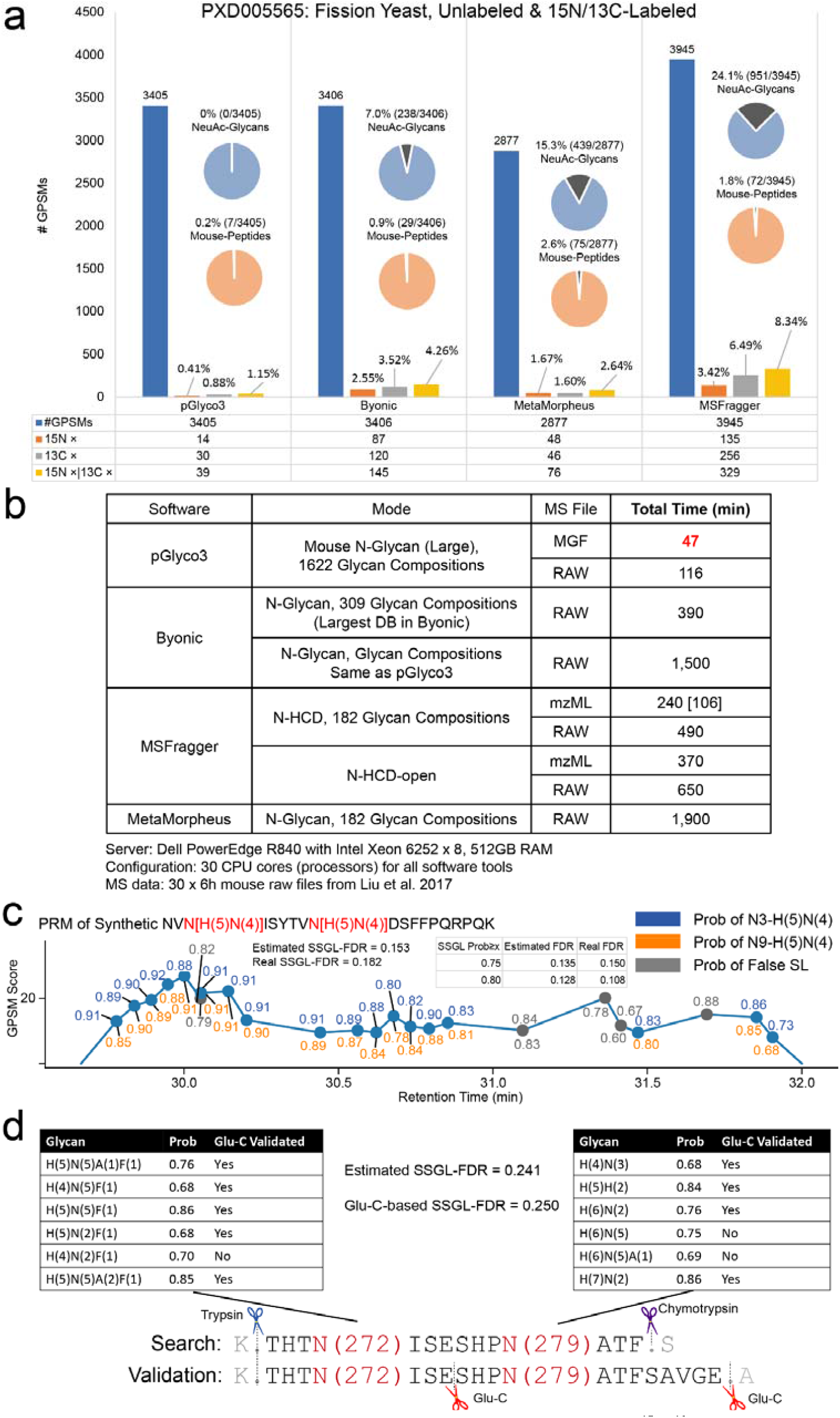
Analyzing N-glycopeptides using pGlyco3. (a) Accuracy comparison using ^15^N/^13^C-labeled fission yeast data (PXD005565). The element-level error rate (incorrect number of N or C elements) of the identified glycopeptides was tested via the ^15^N/^13^C-labeled precursor signals. The potential glycan-level error rate was tested by the percentage of NeuAc-containing GPSMs, and the potential peptide-level error rate was tested via GPSMs with mouse peptides. pGlyco3 shows the best accuracies at all three levels. (b) Search speed comparison under N-glycopeptide data of five mouse tissues (Liu et al. 2017, PXD005411, PXD005412, PXD005413, PXD005553, and PXD005555, 6 h gradient for each RAW file, 30 RAW files in total). The algorithm part (searching MGF files) of pGlyco3 is 5-40 times faster than that of the other tools, even when using larger glycan databases. MSFragger kernel takes only 106 minutes, but its postprocessing modules use too much time. (c) Validation of pGlycoSite using the PRM of the synthetic double-site N-glycopeptide “NVN[H(5)N(4)]ISYTVN[H(5)N(4)]DSFFPQRPQK”. (d) Validation of pGlycoSite using double-site N-glycopeptides “K.THTN(272)ISESHPN(279)ATF.S” of IGHM digested with chymotrypsin and trypsin. The SSGL results were validated by using Glu-C and trypsin digestion with PRM. The example spectra are annotated in Supplementary Fig. 10. All annotated GPSMs for these 12 localized glycans are displayed in Extended Figures 2 in Supplementary Data 3.

**Figure 3.**
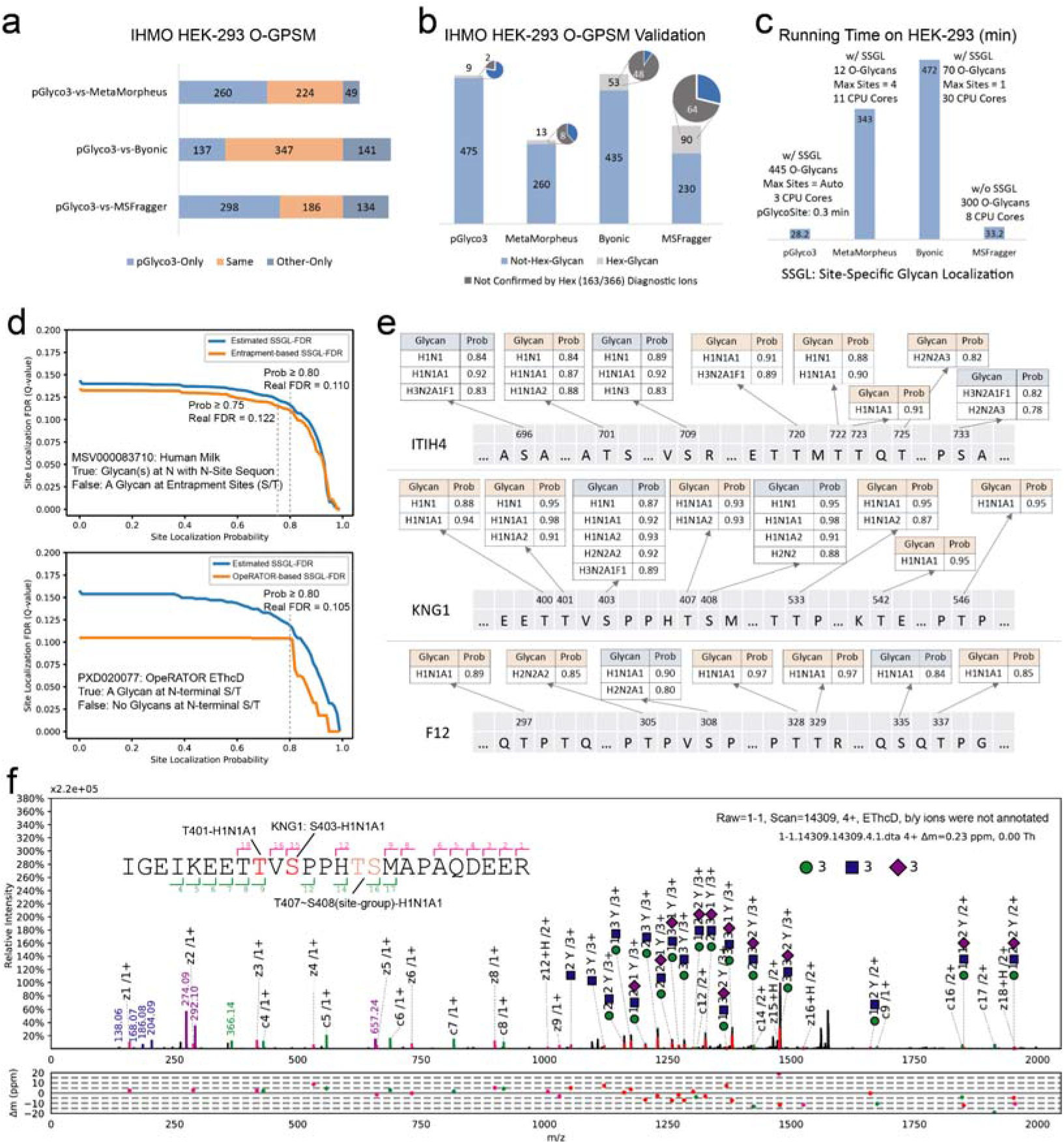
O-glycopeptide identification and SSGL of pGlyco3. (a-c) Software comparisons of O-glycopeptide searches with IHMO HEK-293 cell line data. (a) The overlaps of other tools with pGlyco3 on O-GPSMs. For glycans, only the total glycan compositions were compared; SSGL was not considered. ETD scans in the results of Byonic were mapped to their corresponding HCD scans for the comparisons. If an HCD spectrum and its sister ETD spectrum were identified as the same glycopeptide, we kept only one GPSM; otherwise, we kept both. (b) The identified IHMO O-GPSMs were validated by Hex-containing results and further validated by Hex-diagnostic ions. (c) The runtime comparison. (d) Validation of the SSGL-FDR of pGlycoSite using the entrapment-based SSGL-FDR and OpeRATOR-based SSGL-FDR (Online Methods). (e) Localized site-specific O-glycans of ITIH4, KNG1, and F12 proteins in human serum samples. Site-groups were discarded and SSGL assignments with maximal probability ≥0.75 were displayed. (f) An annotated spectrum of the localized O-glycan and its SSGL probability for KNG1-S403. The HCD spectrum is annotated in Supplementary Fig. 11. The *ScoreTable* with *BestPath* is displayed in Extended Figures 1 in Supplementary Data 3.

Next, we compared the runtime of pGlyco3 with that of other software tools on large-scale mouse N-glycopeptide data from our previous work^8^ (Supplementary Data 1). All these tools including pGlyco3, use multiprocessors to accelerate the searches. We used 30 processors for all the tools to search the data on a Dell workstation with 64 CPU cores and 512 GB physical memories. The time comparisons are shown in Fig. 2b. Ignoring the RAW file parsing time (searching from MGF files), pGlyco3 took only 47 minutes to finish the search (∼1.6 minutes per file, 1622 glycan compositions in the database). The second-fastest tool, MSFragger, took 240 minutes (∼8 minutes per file; 106 minutes for MSFragger kernel excluding its postprocessing modules, 3.53 minutes per file) with an ∼7 times smaller glycan database (182 glycan compositions). The running time of pGlyco3 starting from RAW files was ∼3.9 minutes per file, which was also faster than that of the other tools. The runtime comparisons provided strong evidence showing that the glycan-first search with glycan ion-indexing is very fast for glycopeptide identification.

Finally, we validated pGlycoSite on benchmarked double-site N-glycopeptides. We first synthesized an N-glycopeptide “NVN[H(5)N(4)]ISYTVN[H(5)N(4)]DSFFPQRPQK”, and used parallel reaction monitoring (PRM) to trigger its HCD and EThcD spectra. After identification and localization, we compared the SSGL results with synthetic templates to evaluate the accuracy of SSGL-FDR. As shown in Fig. 2e, among all the SSGL results, the estimated SSGL-FDR was 0.153, which was close to the real SSGL-FDR (0.182), indicating that the SSGL probabilities estimated by pGlycoSite did not have a large deviation. We also used pGlyco3 to identify and localize the double-site N-glycopeptide “K.THTN(272)ISESHPN(279)ATF.S” of IGHM digested from chymotrypsin and trypsin and validated the localized double-site glycans by using Glu-C digestion and PRM (Fig. 2f). pGlyco3 localized 12 site-specific N-glycans on N272 and N279, with an estimated SSGL-FDR 0.241. Nine of 12 N-glycans could be confirmed by further Glu-C digestion, showing a Glu-C-suggested SSGL-FDR of 0.250.

Overall, based-on the glycan-first search and glycan-level quality control, pGlyco3 outperformed the other three software tools in terms of accuracy and search speed. Validation results on double-site N-glycopeptides showed that SSGL probabilities estimated by pGlycoSite were quite accurate.

### pGlyco3 for O-glycopeptide identification

As the glycan-first search of pGlyco3 has been shown to be accurate and fast for N-glycopeptide identification, we then demonstrated the performance of pGlyco3 for O-glycopeptide identification. The workflow of O-glycopeptide is similar to that of N-glycopeptide, except that S/T are the candidate glycosylation sites, and the core Y ions are changed (Supplementary Table 2). We compared O-glycopeptide identification with pGlyco3 to that with tools using inhibitor-initiated homogenous mucin-type O-glycosylation (IHMO) cell line datasets (Fig. 3a-c, Supplementary Note 4). IHMO cells were almost inhibited with only truncated HexNAc(1) or HexNAc(1)NeuAc(1) O-glycans on the peptides (Supplementary Note 4). The identified O-GPSMs on the IHMO HEK-293 dataset (Fig. 3a) were then evaluated by Hex-containing GPSMs, which were further confirmed by checking the Hex-diagnostic ions (163.060 and 366.139 m/z, Fig. 3b, Online Methods). pGlyco3 obtained only 1.9% (9 of 484) of Hex-GPSMs, and only 2 of 9 Hex-GPSMs could not be validated by the Hex-diagnostic ions, showing a very low Hex-suggested glycan-level error rate (2/484≈0.4%) for the IHMO HEK-293 data. The Hex-suggested glycan-level error rates of the three other software tools on the same dataset were: ∼2.9% (8/273) for MetaMorpheus, ∼10.0% (48/488) for Byonic, and ∼20.0% (64/320) for MSFragger. These results further proved the accuracy of pGlyco3 in glycopeptide identification. The search speed of pGlyco3 was also higher than that of others on the same IHMO dataset.

To demonstrate the accuracy of pGlycoSite for SSGL on O-glycopeptides, we designed two experiments to validate the estimated SSGL-FDR: entrapment-based and OpeRATOR-based methods. The entrapment-based method applies pGlycoSite on N-GPSMs by regarding J/S/T (J is N with the N-glycosylation sequon) as the candidate sites, and SSGL would be false in an N-GPSM if a site is not localized at J. The OpeRATOR-based method suggests that SSGL would be false for a GPSM if no glycans are localized at the N-terminal S/T since the OpeRATOR recognizes O-glycans and cleaves O-glycopeptides at the N-termini of O-glycan-occupied S or T^36^. The entrapment-based SSGL-FDR and OpeRATOR-based SSGL-FDR could be deduced (Online Methods). SSGL-FDRs estimated by pGlycoSite were validated by entrapment-based SSGL-FDR and OpeRATOR-based SSGL-FDR, and the results showed that SSGL-FDRs estimated by pGlycoSite did not have much deviation, as shown in Fig. 3d and Supplementary Fig. 2. Coupling with the validation results of double-site N-glycopeptides (Fig. 2e-f), the entrapment-based SSGL-FDR and OpeRATOR-based SSGL-FDR (Fig. 3d) could verify the accuracies of pGlycoSite for SSGL on both N- and O-glycopeptides. Fig. 2c and 3d also suggested that, the real SSGL-FDR would be nearly 10% at SSGL probability ≥0.8, and the SSGL-FDR would be also acceptable at SSGL probability ≥0.75 (“level-1” SSGL in MetaMorpheus O-pair). Therefore, we recommend 0.8 or 0.75 as the SSGL probability cut-off value, but one should know that the real SSGL-FDRs may be different for different datasets at the same SSGL probability threshold.

Finally, we used pGlycoSite to analyze O-glycopeptides in human serum samples, and the localized site-specific O-glycans with probabilities for the proteins inter-alpha-trypsin inhibitor heavy chain H4 (ITIH4), kininogen-1 (KNG1), and coagulation factor XII (F12) are shown in Fig. 3e. All these localized O-glycosylation sites except for S403 on KNG1 can be confirmed on www.uniprot.org or www.oglyp.org^37^, showing the reliability of these localized sites. The O-glycosylation of S403 on KNG1 can be verified by EThcD data, as shown in Fig. 3f. We also displayed the zoom-in regions of the key c/z ions for SSGL of this GPSM as well as a few other GPSMs in Extended Figure 2 in Supplementary Data 3. Based on this dataset, inspired by the work of MetaMorpheus O-pair^21^, we also evaluated the search times and peptide FDRs using different entrapment protein databases (Supplementary Fig. 9). Results showed that pGlyco3 is fast and accurate even with large number of entrapment peptide sequences.

### Analyses of aH-glycopeptides in yeast samples

In the previous sections, pGlyco3 was proven to be reliable for glycopeptide identification, we then applied pGlyco3 to analyze aH-glycopeptides in yeast samples. In this manuscript, “aH” refers to the Hex with an ammonium adduct^38^.

We first searched N-glycopeptides and O-Man glycopeptides in the ^15^N/^13^C labeled fission yeast dataset (PXD005565^8^) with Hex “modified” by aH (Online Methods). Then, the identified aH-glycopeptides were confirmed at the element level by ^15^N- and ^13^C-labeled precursors. As shown in Fig. 4a, we found 579 unique aH-N-glycopeptides with one aH and 164 aH-N-glycopeptides with two aHs. In total, 571 of 579 and 155 of 164 aH-glycopeptides could be confirmed by the ^15^N and ^13^C MS1 evidence for aH×1 and aH×2, respectively. 86% (22 + 21 of 50) of aH-N-glycans identified in glycopeptides could be confirmed by MS/MS data of released N-glycans, as shown in Fig. 4a. aH could also be identified on O-Man glycopeptides in PXD005565 and confirmed by ^15^N and ^13^C MS1 evidence (Fig. 4a, right).

**Figure 4.**
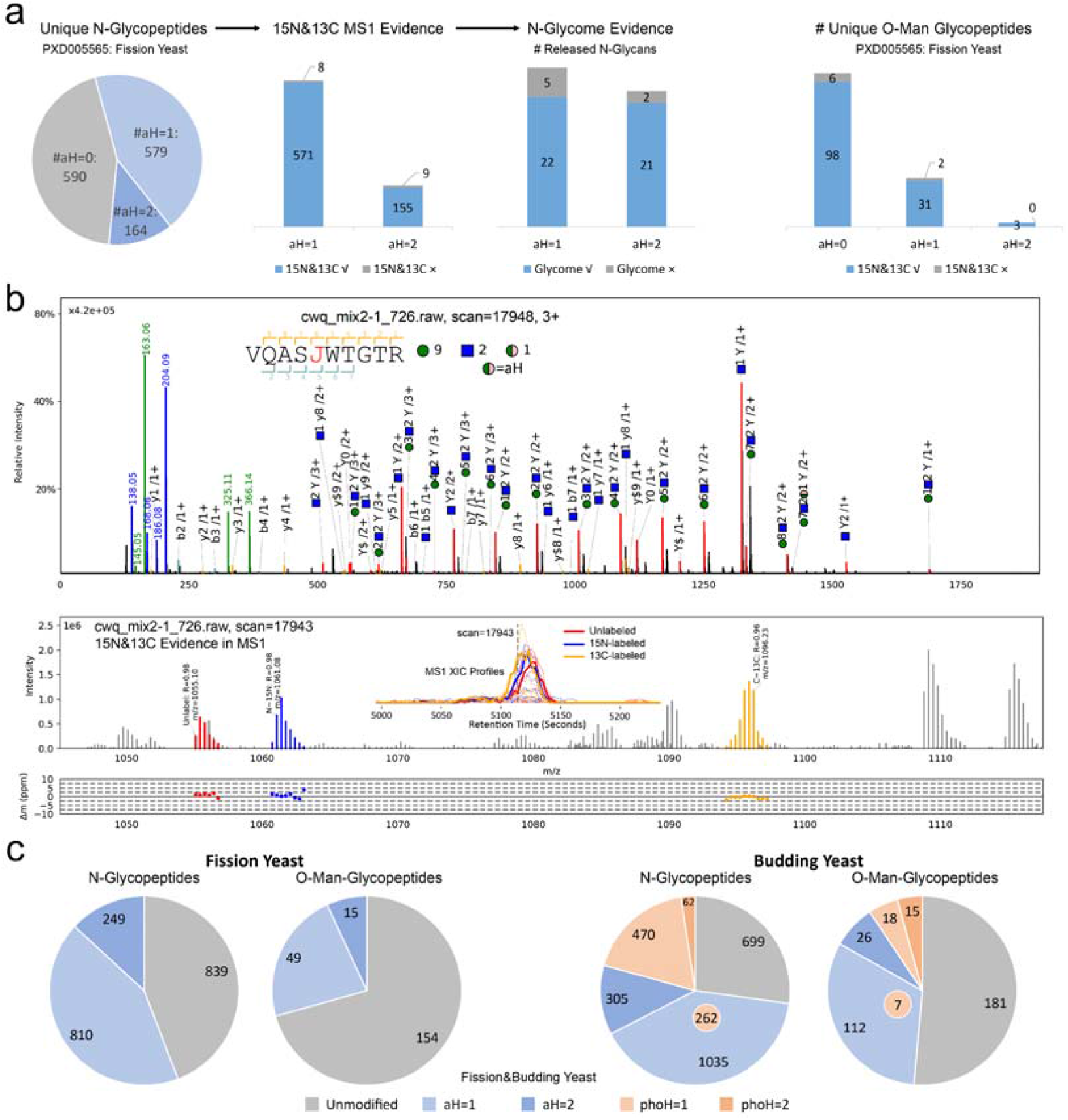
Analysis and verification of the “Hex+17 (aH)”. (a) Unlabeled aH-N-glycopeptides are identified on fission yeast data (PXD005565) and are verified by the ^15^N/^13^C-labeled MS1 precursors and the N-glycome MS/MS data. Unlabeled aH-O-Man glycopeptides are also identified and verified by ^15^N/^13^C-labeled MS1 evidence. (b) MS/MS spectral annotation (top) and ^15^N&^13^C MS1 evidence (bottom). In the MS1 annotation, “R” is the Pearson correlation coefficient between the theoretical and experimental isotope distributions. The blue square and green circle refer to HexNAc and Hex, respectively. The MS2 spectrum is a chimera spectrum of “VQASJ(N)WTGTR + Hex(9)HexNAc(12)aH(1)” and 15N-labeled “VQASJ(N)WTGTR + Hex(10)HexNAc(2)”, see Extended Figure 2 in Supplementary Data 3 for details. (c) Deep analysis of aH-N-glycopeptides and aH-O-Man glycopeptides in fission yeast and budding yeast data. Phospho-Hex (phoH) glycopeptides are also found in budding yeast samples. 262 N-glycopeptides and 7 O-Man glycopeptides in budding yeast contain both aH and phoH.

Fig. 4b and Supplementary Fig. 3d showed the MS/MS, ^15^N and ^13^C MS1, and N-glycome evidence of the aH-N-glycopeptide “VQASJ(N)WTGTR + Hex(9)HexNAc(2)aH(1)”. This glycopeptide was identified at MS/MS scan 17948 and was validated by its unlabeled, ^15^N-labeled, and ^13^C-labeled MS1 precursors at scan 17943. The MS/MS annotation showed that the aH-glycan was confidently identified with many continuous Y ions matched (Fig. 4b, top). MS1 precursor evidence showed high Pearson correlation coefficients (Rs) and low matching mass errors for all the unlabeled, ^15^N-labeled and ^13^C labeled precursors (Fig. 4b bottom).

Fig. 4a and 4b showed that pGlyco3 can confidently identify aH-glycopeptides. We then generated large-scale fission yeast and budding yeast datasets to further analyze the aH-glycopeptides (Supplementary Note 4, the search parameters are shown in Supplementary Data 1), and the results are shown in Fig. 4c. We found large proportions of aH-glycopeptides in both fission yeast and budding yeast samples. In total, 55.8% (1059 of 1898) of aH-N-glycopeptides and 29.4% (64 of 218) of aH-O-Man glycopeptides were identified in fission yeast. The percentages were 58.0% and 40% in budding yeast. pGlyco3 also identified several phoH-N-glycopeptides and phoH-O-Man glycopeptides in the budding yeast dataset. We analyzed site-specific N-glycans and O-Man glycans of protein O-mannosyltransferase (ogm1) in fission yeast, and the O-Man sites were localized by pGlycoSite on EThcD data, as displayed in Supplementary Fig. 5. aH-glycopeptides were also found in other public glycopeptide datasets (Supplementary Fig. 8), demonstrating the commonality and importance of aH.

## Discussion

In this work, we emphasized the importance of glycan-level quality control for the development of glycopeptide search engines to ensure the glycan-level accuracy. A glycan is not just a simple PTM, its mass is sometimes so large that traditional PTM searches may obtain different glycan compositions or even different glycan and amino acid combinations^23^. Unexpected peptide-level modifications could also lead to incorrect glycan identfications^39^. Fortunately, glycans have their own fragment ions and diagnostic ions, allowing search engines to achieve more accurate glycan identification. Strict glycan-level quality control may reduce the sensitivity, but accuracy should always be the first priority for any search engine. Therefore, for peptide-first search engines, we also recommend performing glycan-level quality control after the peptides are identified, as they have obtained superior peptide identification performance.

SSGL is important, especially for O-glycosylation. MetaMorpheus uses a graph-based algorithm to localize sites and estimate site probabilities, but it needs to build different graphs for different combinations of glycans. pGlycoSite provides a one-step algorithm to deduce the sites based on the *ScoreTable* with a dynamic programming algorithm, making the SSGL step extremely fast. As heuristic algorithms for glycosylation SSGL have been developed only in the last few years, there is still plenty of room to improve the scoring schema, SSGL probability estimation, and result validation.

pGlyco3 provides a convenient way to analyze modified saccharide units based on canonicalization-based glycan representations, making the analysis of modified saccharide units similar to that of peptide modifications for users. This new feature enables us to identify aH-glycopeptides, which could be validated through N-glycome data and ^15^N/^13^C-labeled data, on N-glycopeptides and O-Man glycopeptides in yeast samples. aH may be not easy to avoid as it could also be found in many other public glycopeptide datasets.

In Supplementary Note 5, based on the PXD005565 yeast dataset, we further discussed how different glycan databases influenced the identification; why Byonic obtained different glycopeptides; and more importantly, how to improve our glycopeptide identifications. The results also showed the pros and cons of searching for aH during glycopeptide identification; therefore, the identification interferences caused by aH should be considered in the routine glycoproteomic analyses. We also found some common errors for different search engines (e.g., precursor detection error), suggesting possible future improvements for all software tools including pGlyco3.

## Online Methods

### Spectrum preprocessing

pGlyco3 can process HCD and “HCD+ETxxD” (HCD-pd-ETxxD, or HCD followed by ETxxD) data for N-/O-glycopeptide identification; here, ETxxD could be either ETD, EThcD, or ETciD. Note that pGlyco3 is optimized for both glycan and peptide fragment analysis using sceHCD; hence, sceHCD is always recommended, as discussed in our previous work^8^. If HCD and ETxxD spectra are generated for the same precursor, pGlyco3 will automatically merge them into a single spectrum for searching. Peaks from HCD and EThcD spectra are merged based on a specified mass tolerance set by the users (±20 ppm by default) in the spectrum preprocessing step. pGlyco3 then deisotopes and deconvolutes all MS2 spectra and removes the precursor ions. pGlyco3 uses pParse to determine the precursor mono ions and to export chimera spectra. The pParse module has been also used in peptide, glycopeptide and cross-linked peptide identification^40, 41^. pGlyco3 filters out non-glycopeptide spectra by checking glycopeptide-diagnostic ions, then searches the glycan parts with the glycan ion-indexing technique. The diagnostic ions can be defined by the users, the default ion is 204.087 m/z for both N-glycosylation and O-glycosylation.

### Glycan-first search and glycan ion-indexing

In pGlyco3, each glycan structure is represented by a canonical string in the glycan database. pGlyco3 provides quite a few built-in N- and O-glycan databases, and it supports the conversion of the glycan structures of GlycoWorkbench^35^ into the canonical strings of pGlyco3. For modified saccharide units, pGlyco3 will automatically substitute one or several unmodified monosaccharides in each canonical string into modified forms, making it very convenient for the modified saccharide unit analysis (Supplementary Note 1).

For each peak in an MS2 spectrum, pGlyco3 assumes it is a Y ion of a glycopeptide with the peptide attached to the database search. In pGlyco 2.0, we calculated the peptide mass for each glycan in the glycan database by “precursor mass – glycan mass”, and theoretical Y ion masses of each glycan could be deduced and matched against the MS2 spectrum. This glycan search algorithm has to match the same spectrum for N times (O(N), N is the number of glycans in the glycan database). But pGlyco3 searches for Y-complementary ions (“precursor mass – Y ion mass”) instead of Y ions, the key observation was that:

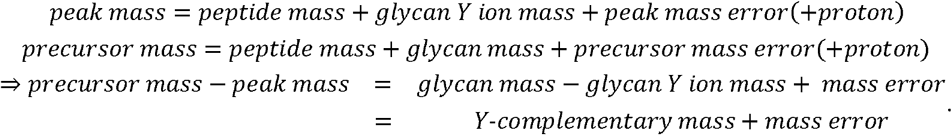

To accelerate the Y-complementary ion search, pGlyco3 builds the ion-indexing table for Y-complementary ions of all possible Y ions for the glycan database (Supplementary Note 2). The Y-complementary ion composition is defined as “full glycan composition – Y-ion glycan composition”. A table of all possible unique Y-complementary ion compositions is generated, and the list of glycan IDs where the Y-complementary compositions originate is also recorded in the table. For the glycan-first search, core Y ions (e.g., trimannosyl core Y ions in N-glycans, Supplementary Table 2) are the key ions for N-glycan scoring. Hence, pGlyco3 encodes the glycan ID using 31 bits of the 32-bit integer and uses the extra bit to record whether the corresponding Y ion is the core ion of the glycan (Supplementary Note 2). The table is then sorted and hashed by the masses for a fast query (within the O(1) query time). As a result, it takes only O(#Peak) time to obtain the matched ion counting scores as well as the matched core ion counting scores of all glycans for every spectrum; here, #Peak refers to the number of peaks in a spectrum. pGlyco3 retains the glycans with ≥ n core Y ions matched (n = 2 for an N-glycan and n = 1 for an O-glycan). Glycans are further filtered out if they contain a specific monosaccharide but are not supported by corresponding monosaccharide-diagnostic ions (Supplementary Table 1). Finally, pGlyco3 retains the top 100 candidate glycan compositions (sum of the ion counting score and the core ion counting score) for the peptide search. pGlyco3 also retains all small glycans (number of monosaccharides ≤ 3) for the peptide search because small glycans have too few Y ions to obtain high glycan scores.

### Peptide search and glycopeptide FDR estimation

In protein sequence processing, every Asn (N) with the sequon “N-X-S/T/C (X is not P)” in all protein sequences is converted to “J” while keeping the same mass and chemical elements as N. J is then the candidate N-glycosylation site, and S/T are the candidate O-glycosylation sites. Proteins are digested into peptide sequences, and peptide modifications are added to the peptide sequences. Modified peptides are also indexed by their masses for O(1) time access. For the given spectrum and each candidate glycan, the peptide mass is deduced by “precursor mass – glycan mass”. pGlyco3 then queries the peptide mass from the mass-indexed peptides. For the peptide search, pGlyco3 considers b/y ions and “b/y + HexNAc” in the HCD mode. pGlyco3 further considers c/z ions as well as their hydrogen rearrangement in the merged spectra for the HCD+ETxxD mode. The candidate glycan is also fine-scored by the matched Y ions. The glycan and peptide scoring schemes of pGlyco3 are the same as those of pGlyco 2.0, but some parameters were tuned in pGlyco3 to obtain better identification performance (Supplementary Fig. 6). Only the top-ranked glycopeptide is retained as the final result for each spectrum. For potential chimeric spectra, pGlyco3 removes unreliable mixed glycopeptides by determining whether one’s precursor is another’s isotope. For example, if NeuAc(1) and Fuc(2) are simultaneously identified in the same MS2 scan but with different precursors, the Fuc(2)-glycopeptide will be removed because “NeuAc(1) + 1 Da = Fuc(2)”. pGlyco3 also uses pGlyco 2.0’s method to estimate the FDRs for all GPSMs at the glycan, peptide, and glycopeptide levels. pGlyco3 skips the glycan FDR estimation step for small glycans.

### Fast glycosylation site localization with pGlycoSite

pGlyco3 determines not only the glycosylation sites but also determines the attached glycan composition to each site. For a given spectrum with the identified peptide and glycan composition, enumerating all possible glycopeptide-forms and generating their c/z ions for site localization is not computationally easy. The worst computation complexity could be 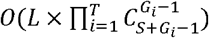, where *L* is the peptide length, *T* is the number of monosaccharide types, *S* is the number of candidate glycosylation sites, and *G*_*i*_ is the number of the *ith* monosaccharide type (Supplementary Note 3). The enumeration complexity would be exponentially large as *S* and *G*_*i*_ increase, as illustrated in Supplementary Note 3.

In pGlyco3, the pGlycoSite algorithm is designed to avoid enumeration. The key observation for the pGlycoSite algorithm is, that regardless of how many glycopeptide forms there are for a given peptide and glycan composition, the number of all possible c or z ions is at most *F* × (*L* − 1). Here, *F* is the number of subglycan compositions for the identified glycan composition, and the subglycan is defined in Supplementary Note 3. pGlyco3 generates a c/z-ion table in which each cell contains the glycan-attached c/z ions (Supplementary Note 3). After removing all Y and b/y ions from the given spectrum, the ion table with c/z ions is then matched and scored against the spectrum (called the *ScoreTable*, as illustrated in Fig. 3b and Supplementary Note 3). pGlycoSite currently uses c/z ion counting scores for each cell of the table, but other comprehensive scoring schemes could be supported in the table if they could achieve better performance.

The best-scored path starting from bottom left [*g*_0_,0] to top right [*G,L*] (Fig. 3c and Supplementary Note 3) is then calculated by a dynamic programming algorithm:

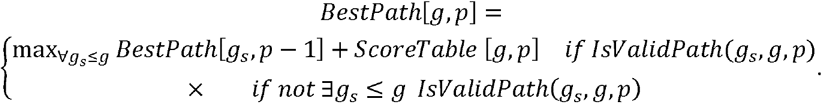

where *G* is the identified full glycan composition, *g* refers to a subglycan composition of *G, g*_0_ refers to the zero-glycan composition, and *p* refers to *p*th position of the peptide sequence. Here, all glycan compositions (from *g*_0_ to *G*) are represented as vectors and hence could be compared with each other. Therefore, *g*_*s*_ ≤ *g* means that *g*_*s*_ is the subglycan of *g. IsValidPath* (*g*_*S*_, *g, p*) is designed to check whether the path starting from [*g*_*s*_,p−1] to [*g,p*] is valid (Supplementary Note 3). pGlycoSite sets *BestPath*[*g*,0] = 0 (∀*g*: *g*_0_ ≤ *g* ≤ *G*) and iteratively calculates the *BestPath* table for all *g*_0_ ≤ *g ≤ G* and 0 < *p* ≤ *L*. *BestPath*[*G,L*] is then the final best path score that will be solved. Finally, pGlycoSite deduces all the paths that can reach the *BestPath*[*G,L*] score by backtracking the *BestPath* table from [*g*_0_,0] to [*G,L*]. If the best-scored path contains the cell [*g*_*s*_,*p* − 1] and [*g,p*] with *g*_*s*_ < *g*, then the *p*th amino acid is localized as a site with glycan *g* − *g*_*s*_. pGlycoSite introduces the “site-group” if multiple paths can achieve the same *BestPath*[*G,L*] score (Fig. 3c, Supplementary Note 3). The time complexity of SSGL in pGlycoSite including the dynamic programming and backtracking for a GPSM, is only *0*(*L* × *F*^2^) (Supplementary Note 3).

### Site localization probability estimation with pGlycoSite

Glycosylation site probability refers to the probability that a site is correctly localized. As the peptide and glycan compositions have been identified for a given MS2 spectrum, an incorrect localization would result from the random assignment of randomly selected subglycans to random sites for the same peptide and glycan compositions. To simulate the incorrect localization for each localized site, pGlycoSite randomly samples 1000 paths from bottom left to top right on the *ScoreTable*. For a given site or site-group to be estimated, the random paths could overlap with the *BestPath*, but they must not contain the path that can determine this site or site-group (i.e., path from [*g*_*s*_,*p*_*i*_] to [*g,p*_*j*_] for site *p*_i_ (*j* = *i* +1) or site-group {*p*_*i*_,*p*_*j* − 1_} (*j* > *i* + 1)). pGlycoSite then calculates 1000 ion counting scores of these paths, and estimates a Poisson distribution from these random scores. It estimates the *p*-values based on the Poisson distribution for the *BestPath*[*G,L*] and the best random score (denoted as *RandomBest*) and then estimates the probability as follows:

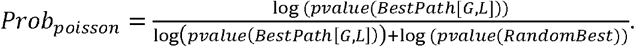

To ensure the localized glycopeptide-spectrum-matching quality, pGlycoSite adds a regularization factor to the estimated *Prob*_*poisson*_, and the final localized probability becomes

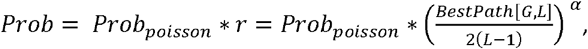

where *α* is set as a small value (0.05) to ensure that it does not affect the value of *Prob*_*poisson·*_.

However, when *BestPath*[*G,L*] obtains a very small score, *r* will be close to zero, hence limiting the final *Prob* value. *L*−1 is the number of the considered c/z ions.

### Benchmark, software versions

pGlyco3 was compared with MetaMorpheus (v0.0.312, downloaded in 2020.10), MSFragger (v3.1.1 with FragPipe v14.0 and philosopher v3.3.11, downloaded in 2020.10), and Byonic (v3.10).

### Previously published mass spectrometry data

sceHCD RAW files of mixed unlabeled, ^15^N-labeled, and ^13^C-labeled fission yeast glycopeptide samples were downloaded from PXD005565^8^ on PRIDE. Moreover, 30×6 h sceHCD RAW files of five mouse tissues were downloaded from PXD005411, PXD005412, PXD005413, PXD005553, and PXD005555^8^ on PRIDE. sceHCD-pd-EThcD RAW files of human milk and Chinese hamster ovary cell (CHO) samples were obtained from MassIVE (dataset MSV000083710^7^). RAW files of OpeRATOR-processed O-glycopeptide data were obtained from PXD020077^10^ on PRIDE. Detailed search parameters for all these RAW data files are listed in Supplementary Data 1.

### Validation of the N-glycopeptide search with ^15^N/^13^C-labeled fission yeast data

The protein sequence database used was the fission yeast protein sequence database (S. pombe, Swiss-Prot, 2018.08) concatenated with the mouse protein sequence database (M. musculus, Swiss-Prot, 2018.08). Identified GPSMs with mouse peptides would be falsely identified, and hence, mouse peptide GPSMs could be used to test the peptide-level error rates. The N-glycan database for MetaMorpheus, MSFragger, and Byonic is the 182-glycan database which includes 74 NeuAc-containing N-glycan compositions. The N-glycan database for pGlyco3 is the built-in mouse N-glycan database, which contains 1234 N-glycan compositions (6662 structures) and has 659 NeuAc-contained compositions. NeuAc-containing N-glycan compositions identified in fission yeast data would be falsely identified and thus could be used to test the glycan-level error rates. The detailed search parameters are listed in Supplementary Data 1. For each software tool, all spectra were regarded as unlabeled spectra while searching, and the identified GPSMs were then validated by using their ^15^N/^13^C-labeled precursor signals in the MS1 spectra (Fig. 2a). This validation method was also used in our previous works for peptide, glycopeptide, and crosslinked peptide identification^8, 40, 41^. Peptide-level and glycan-level FDRs were also tested by using mouse peptides and NeuAc-containing glycans, respectively (Fig. 2a).

### Validation of the O-glycopeptide search with IHMO data

In Inhibitor-initiated Homogenous Mucin-type O-glycosylation (IHMO), an O-glycan elongation inhibitor, benzyl-N-acetyl-galactosaminide (GalNAc-O-bn), was applied to truncate the O-glycan elongation pathway during cell culture, generating cells with only truncated HexNAc(1) or HexNAc(1)NeuAc(1) O-glycans. sceHCD-pd-EThcD spectra were generated after O-glycopeptides were enriched by FASP^42^, and experimental details are shown in Supplementary Note 4. IHMO in HEK-293 cells was then verified by laser confocal microscopy, as displayed in Supplementary Note Fig. 4. Spectra were then searched by pGlyco3, MetaMorpheus, MSFragger, and Byonic, and the search parameters are listed in Supplementary Data 1. For all software tools, Hex-containing O-glycopeptide could be still identified due to the inhibitor’s imperfect efficiencies. The Hex-contained O-GPSMs were further validated by the summed intensities of the Hex-diagnostic ions (163.060 and 366.139 m/z) in their HCD spectra. The summed intensity threshold was set as 10% to the base peak.

### Validation of the pGlycoSite algorithm

For the given SSGL probabilities (*Prob*) of all identified GPSMs, the SSGL-FDR could be estimated as follows:

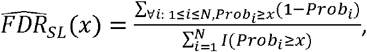

where 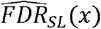 is the estimated SSGL-FDR for a given probability threshold *x, N* is the total number of localized sites, and *I*(*bool*) is the indicator function which returns 1 when *bool* is true and 0 otherwise. It is not easy to validate the estimated SSGL-probability for a given site, but we can validate the accuracy of 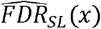, enabling SSGL-probability validation from another perspective. In this work, we designed four methods to validate 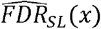, synthetic double-site N-glycopeptide validation, multienzyme-based validation, entrapment-based validation and OpeRATOR-based validation.

A double-site N-glycopeptide “NVN[H(5)N(4)]ISYTVN[H(5)N(4)]DSFFPQRPQK” was synthesized and its 3+ and 4+ HCD+ETxxD spectra were continuously acquired by using PRM. To enable SSGL validation, we searched the spectra against the human glycan database and a protein database with only the synthetic peptide sequence. Thus, we could always identify the same peptide sequence with different localized glycans. SSGL would be true if the localized glycan was H(5)N(4); otherwise, it was false. The real SSGL-FDR could then be calculated as 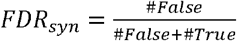.

To validate SSGL for double-site N-glycopeptides under more comprehensive situations, we used pGlyco3 to identify and localize the double-site N-glycopeptides with the peptide sequence “K.THTN(272)ISESHPN(279)ATF.S” of IGHM digested with chymotrypsin and trypsin. For all identified site-specific N-glycans, the theoretical masses of further “trypsin+Glu-C”-digested glycopeptides were calculated. Then, we also used PRM to trigger the HCD spectra of “trypsin+Glu-C”-digested glycopeptides and identified these single-site spectra to verify the double-site N-glycopeptides. The SSGL would be true if the localized N-glycan on “K.THTN(272)ISESHPN(279)ATF.S” could be identified by the single-site spectra; otherwise, it was false. The Glu-C-suggested SSGL-FDR could then be calculated as 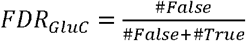.

For entrapment-based validation, after N-glycopeptide data were searched, the sites of GPSMs were localized using pGlycoSite by regarding the candidate sites as “J/S/T”, which could be enabled by setting “glycosylation_sites=JST” in the search parameter file. For a given GPSM, it would be a true positive (*TP*_*trap*_) if J were the only localized sites; otherwise, all sites were false positives (*FP*_*trap*_). The entrapment-based SSGL-FDR could be calculated as

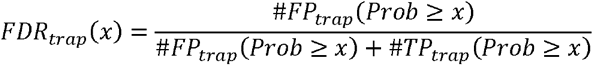

for a given probability threshold *x*. Then, the SSGL probabilities of pGlycoSite could be validated by comparing 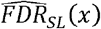 with *FDR*_*trap*_(*x*).

For OpeRATOR-based validation, we used the data digested by OpeRATOR^36^. Only the GPSMs with their peptides starting with ST at the N-terminal were used for the validation. Then, for a given GPSM, we regarded it as the true positive (*TP*_*OpR*_) if localized sites contained a site that was at the N-terminal S/T otherwise, it was regarded as a false positive (*FP*_*OpR*_). The OpeRATOR-based SSGL-FDR (*FDR*_*OpR*_(*x*)) could be calculated from *TP*_*OpR*_(*x*) and *FP*_*OpR*_(*x*) which is similar to *FDR*_*trap*_(*x*).

Comparisons of the 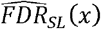 of pGlycoSite with *FDR*_*trap*_(*x*) and *FP*_*OpR*_(*x*) are displayed in Fig. 3d and Supplementary Fig. 2. We also compared 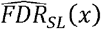 of MetaMorpheus with *FDR*_*OpR*_(*X*), as shown in Supplementary Fig. 2.

### Analysis of aH-glycopeptides in yeast samples

“aH” is defined as a Hex with an ammonia adduct. Peptides were searched by the yeast protein sequence databases (*S. pombe* for fission yeast and *S. cerevisiae* for budding yeast, Swiss-Prot, 2018.08). N-glycan parts were searched against the high-mannose-only N-glycan database, and O-Man glycan parts were searched against the Hex-only glycan database. aH was regarded as a modified Hex for the pGlyco3 search, and the maximal number of aHs was set as 2 per glycan. For the O-Man-glycopeptide search, the glycopeptide-diagnostic ion was set as Hex (163.060 m/z). The ^15^N/^13^C-labeled fission yeast (PXD005565) results were also validated by the ^15^N-labeled and ^13^C-labeled precursor signals in the MS1 spectra. For the ^15^N/^13^C validation of aH identifications, as the ammonia adduct may be introduced during sample processing or MS steps, it could not be labeled by ^15^N; hence we computationally designed a new element called “14N”, which would not be converted into ^15^N for MS1 signal extraction. The element composition is recorded in the “element.ini” file in the software package, demonstrating the flexibility of pGlyco3 for the analysis of new monosaccharides or modified saccharide units. We also verified the aH-N-glycans by analyzing the MS data of released N-glycans in fission yeast samples (Supplementary Note 4).

O-Man glycopeptides of fission yeast were also analyzed by HCD followed by EThcD to investigate the O-mannosylation sites. The data were searched by the “HCD+EThcD” mode of pGlyco3, and the sites were localized by pGlycoSite. See Supplementary Note 4 for details.

## Acknowledgements

We thank Prof. Matthias Mann from Max Planck Institute of Biochemistry for editing this manuscript and the point-by-point replies during the revision. We thank Prof. Wei Huang from Shanghai Institute of Materia Medica, CAS for kindly providing us the synthetic N-glycopeptide samples. This work is supported by grants from the National Key Research and Development Program (2018YFC0910302 to W.-F.Z. / M.-Q.L., 2016YFA0501300 to S.-M.H. / M.-Q.L. / W.-Q.C., and 2016YFB0201702 to P.-Y.Y.), the National Natural Science Foundation of China Project (91853102 to W.-Q.C.), and the innovative research team of high-level local university in Shanghai. This paper is dedicated to the memory of Prof. Peng-Yuan Yang, who passed away during the revision.

## Data Availability

Data generated in this work, including yeast glycoproteomic data, yeast N-glycomic data and IHMO O-glycoproteomic data, could be downloaded from MassIVE with identifier MSV000086771 (username: MSV000086771_reviewer, password: pglyco). All the pGlyco3 result files can be found in Supplementary Data 3.

## Code Availability

pGlyco3 could be downloaded from https://github.com/pFindStudio/pGlyco3/releases. Analysis results and Python Notebooks to reproduce the comparison results could be downloaded from https://figshare.com/projects/Searched_results_and_python_notebooks_for_pGlyco3_manuscript/97592.

## Author Contributions

W.-F.Z. conducted this project, developed the software, and analyzed the data. W.-Q.C. performed the MS experiments and analyzed the data. M.-Q.L. analyzed the data. S.-M.H. and P.-Y.Y. supervised this project. All the authors wrote and revised the manuscript.

## Ethics declarations

Competing Interests

The authors declare no competing interests.

