## Supplementary Information for "Precise, Fast and Comprehensive Analysis of Intact Glycopeptides and Modified Saccharide Units with pGlyco3"

**Supplementary Table 1. Diagnostic ions**

In pGlyco3, the existence of a monosaccharide in a spectrum must be supported by at least one of its diagnostic ions. This is a necessary but not sufficient condition, meaning that a spectrum with the X-diagnostic ions does not have to be identified as an X-containing glycan (X could be any monosaccharide, including NeuAc, NeuGc, Hex, ...). “none” (or “empty”) means pGlyco3 will not check the existence of those monosaccharides by their diagnostic ions.

| Glyco | m/z |
|---|---|
| HexNAc | 204.086648533 |
| Hex | 163.0600994315, 366.1394719645 |
| NeuAc | 292.1026925277, 274.0921278414 |
| NeuGc | 308.0976071498, 290.0870424635 |
| Fuc | (none) |
| aH | (none) |
| phospho-Hex (phoH) | 243.0264304315 |

Users can change these settings in “glyco.ini”. The masses recorded in glyco.ini are the neutral masses. pGlyco3 will automatically calculate the m/z values (as shown in the table) for them. pGlyco3 uses the 204-m/z ion to determine if a spectrum is a glycopeptide spectrum in N-Glycan and O-Glycan mode. It also uses 163-m/z ion to determine if a spectrum is a glycopeptide spectrum for O-mannose (O-Man) glycopeptides. These settings could be modified in the search parameter file. If there is at least one diagnostic ion of monosaccharide X containing >=0.05 intensity relative to the 204-m/z ion, pGlyco3 will not ignore X-containing glycans during the search. Some monosaccharides, for example fucose and aH, often do not have corresponding diagnostic ions in the glycopeptide spectra, hence pGlyco3 does not check their diagnostic ions. In glyco.ini, the users just need to set the diagnostic ions as none (as shown in the table) to disable diagnostic ion checking for these kinds of monosaccharides. In the O-Man mode, the reference ion for comparison is 163 m/z. About how to configure O-Man glycopeptide search, please see Supplementary Table 3 for details. “>=0.05” threshold can be changed in the parameter file.

Other oxonium ions, such as 145.0495347452, 168.0655191604, 186.0760838467, 325.112922863, 657.2348884922, and 673.2298031143, are removed from the spectrum for corresponding monosaccharides and glycans during the glycopeptide fine-scoring step for each glycopeptide candidate to avoid any interferences.

The mass difference between phospho-Hex and sulfate-Hex is very small, making it too difficult to distinguish phospho-Hex from sulfate-Hex using precursor ions in MS1 and diagnostic ions in MS/MS. Therefore, in this work, we did not further determine if a phoH is phospho-Hex or sulfate-Hex. Additional diagnostic ions for sulfoHexNAc, and sulfoHexHexNAc can distinguish sulfated glycopeptides from phosphorylated ones[1]. But in some cases, no diagnostic ions can be observed in the MS2 spectra of sulfated glycopeptides[2]. Therefore, expert knowledge about phosphorylated and sulfated glycopeptides is still needed for the analytes from the user sides. And the post-search analysis for the Y ions and the masses of the diagnostic ions may provide more information to distinguish them.

**Supplementary Table 2. Core Y ions of N- and O-glycan used in this manuscript**

Users can change these settings in the searching parameter file (pGlyco3.cfg) before running pGlyco3. The Y0 ion (peptide moiety only) is always considered as one of the core ions for both N- and O-glycopeptides.

| Type | Core Y-Ion Compositions |
| --- | --- |
| N-Glycan | N(1), N(2), N(2)H(1), N(2)H(2), N(2)H(3), N(1)F(1), N(2)F(1) |
| O-Glycan | N(1), N(2), N(1)H(1), N(1)A(1), N(1)G(1) |
| O-Man | H(1) |

N=HexNAc, H=Hex, F=Fuc, A=NeuAc, G=NeuGc.

**Supplementary Table 3. Key parameters for O-Man glycopeptide identification in yeast samples**

In pGlyco3, the O-Man search could only be done using the command-line interface. Graphical user interface (GUI) only supports common N- and O-glycopeptide search. Users can configure the basic parameters on GUI, then save the parameters (pGlyco3.cfg and multiprocess_run.bat), and then edit and save the parameter file with a text editor, then double click “multiprocess_run.bat” to run pGlyco3.

It is also possible for users to analyze mammalian O-Man glycopeptides[3] and other O-glycosylation types (O-Fuc, O-Glc, etc) using pGlyco3. But users have to define the corresponding glycan databases, core Y-ions, and diagnostic ions for them according to the professional theory knowledge and expert experience.

| pGlyco3.cfg |
| --- |
| ... |
| [glycan] |
| glycan_type=O-Glycan |
| glycan_db=pGlyco-O-HexOnly.gdb |
| glycan_core=**H(1)** |
| gp_marker_glyco=**H** |
| glycan_fix_mod= |
| glycan_var_mod=**H~phoH,H~aH** |
| max_var_mod_on_glycan=2 |
| [protein] |
| ... |

**Supplementary Figure 1. Another example of the pGlycoSite algorithm**

(a) The glycopeptide is “IRTTTSGVPR + HexNAc(4)”, where HexNAc is short for “N”. The sites are localized as “T3:N(1);T4:N(1);T5:N(1);S6:N(1)”. (b) The matched c/z ion table (ScoreTable). (c) Dynamic programming to find the best path score. Red lines indicate the best path. This example as well as Fig. 1d-f are designed for better understanding the pGlycoSite algorithm. This example has only one best path, hence the sites could be uniquely localized (no site-groups). The example in Fig. 1d-f contains branching paths from S3 to T5, so sites from S3 to T5 are regarded as a site-group.

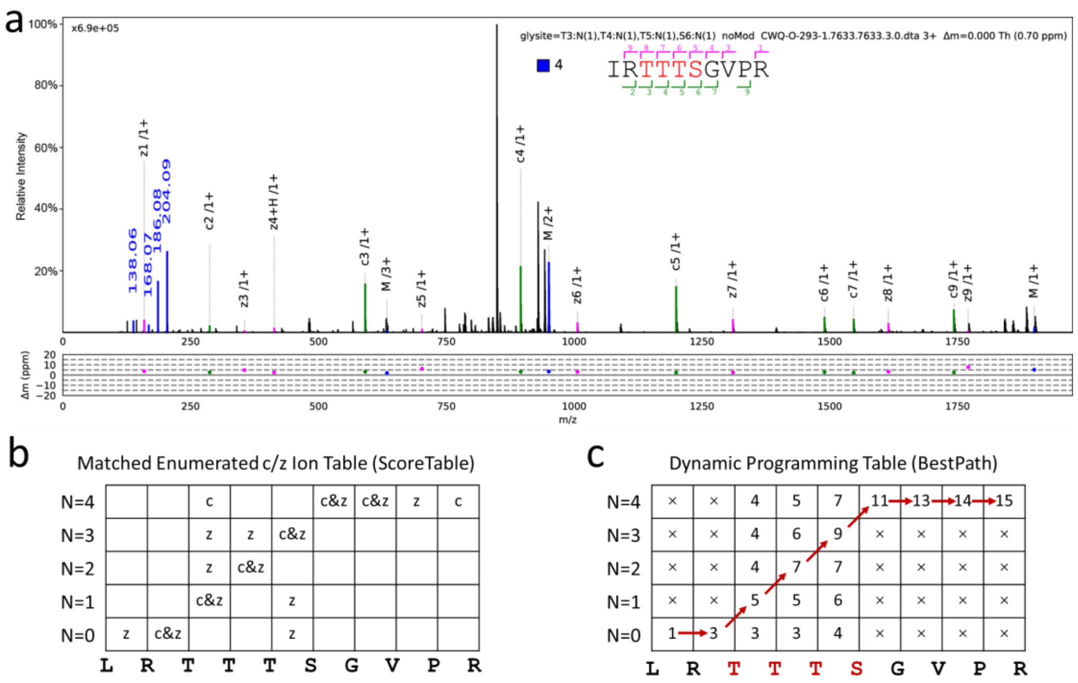

**Supplementary Figure 2. Validation for SSGL-FDR with entrapment-based FDR and OpeRATOR-based FDR**

(a) Validation of SSGL-FDR estimated by SSGL probabilities of pGlycoSite (pGlyco3) using the entrapment-based SSGL-FDR on CHO N-glycopeptide data in the MSV000083710 dataset. (b) Validation of SSGL-FDR estimated by SSGL probabilities of MetaMorpheus using the OpeRATOR-based SSGL-FDR on OpeRATOR O-glycopeptide data in the PXD020077 dataset. Probabilities >= 0.75 are the threshold in MetaMorpheus to determine "level 1" SSGL confidence. This is only to say that the estimated SSGL probabilities of MetaMorpheus may be too optimistic. But with 0.75 cutoff ("level 1"), most of the localized sites tend to be correct. (c) and (d) simulated how the SSGL-FDR curves would change when the digestion specificity (SPC) of OpeRATOR changed (SPC=80%, 90%, and 100%) for OpeRATOR-based validation. Specificity: for all peptide sequences following the OpeRATOR motif, the specificity is defined as the average proportions of GPSMs that contain O-glycosylated N-terminal S/Ts.

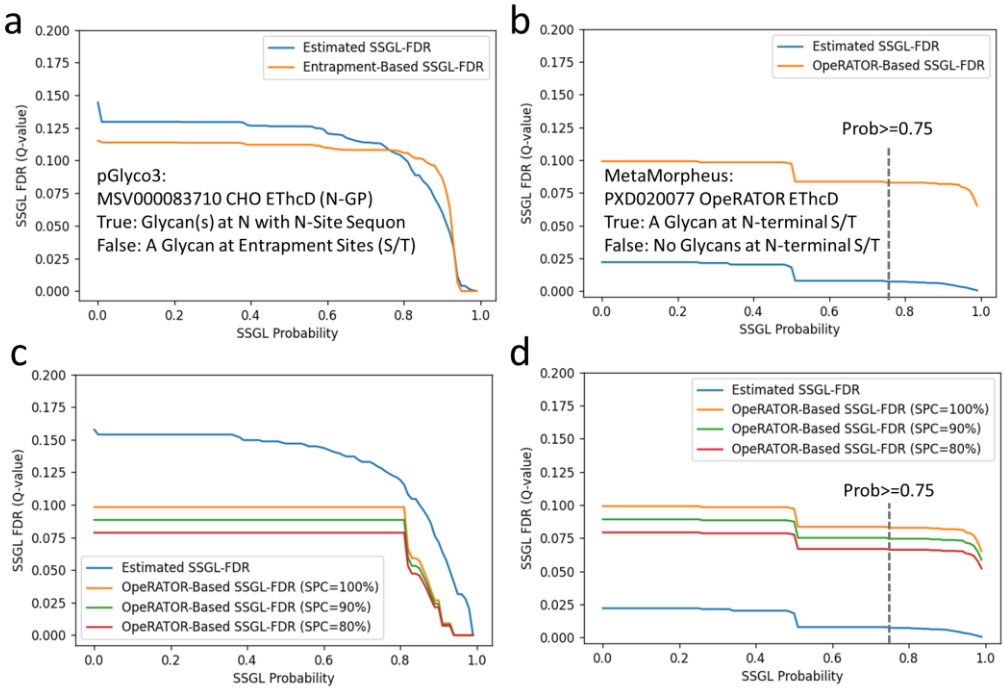

**Supplementary Figure 3. sceHCD-MS/MS, $^{15}N/^{13}C$-labeled MS1, and N-glycome evidences of aH-N-glycopeptides**

The glycopeptide is "YNLJ(N)C[+57]TSE + Hex(11)HexNAc(2)aH(2)" in the $^{15}N/^{13}C$-labeled fission yeast samples. aH is represented by a half-greed-half-pink circle, meaning a modified Hex (green circle). (a) The sceHCD-MS/MS spectrum of the glycopeptide. (b) The preceding MS1 spectrum of the MS/MS spectrum in (a). The unlabeled, $^{15}N$-labeled, and $^{13}C$-labeled precursors and their isotopes are annotated. All these isotope distributions are very similar to their theoretical isotope distributions. R is the calculated Pearson correlation coefficient between experimental and theoretical isotope distributions. The bottom figure in (b) is the extracted isotope patterns through retention time, wherein the colors for unlabeled, $^{15}N$-labeled, and $^{13}C$-labeled peaks are the same as they are at the top figure in (b). (c) N-glycome evidence of "Hex(11)HexNAc(2)aH(2)". These figures proved the existence of aH×2-N-glycopeptides in fission yeast samples. (d) N-glycome evidence of "Hex(9)HexNAc(2)aH(1)" in Fig. 4b in the main text.

(a), (c) and (d) were directly and automatically annotated by gLabel (pGlyco3's built-in GPSM annotation tool) from the original MS/MS spectra (not deisotoped and deconvoluted). (b) was generated by our in-house python script (gLabel-MS1.py). We will integrate gLabel-MS1 into the pGlyco3 system soon to visualize MS1 peaks, or we will support the Skyline PSM format for MS1 visualization. In Supp. Fig. (c) and (d), the B ions are calculated as the sum of the neutral masses of monosaccharides, and the Y ions are calculated as the sum of the neutral masses of monosaccharides plus a water group (18.0105646863 Da). Supp. Fig. (c) and (d) were directly plotted from the RAW file, in which monoisotopic peaks are not determined, hence the $\Delta m$ values are 0.5 m/z.

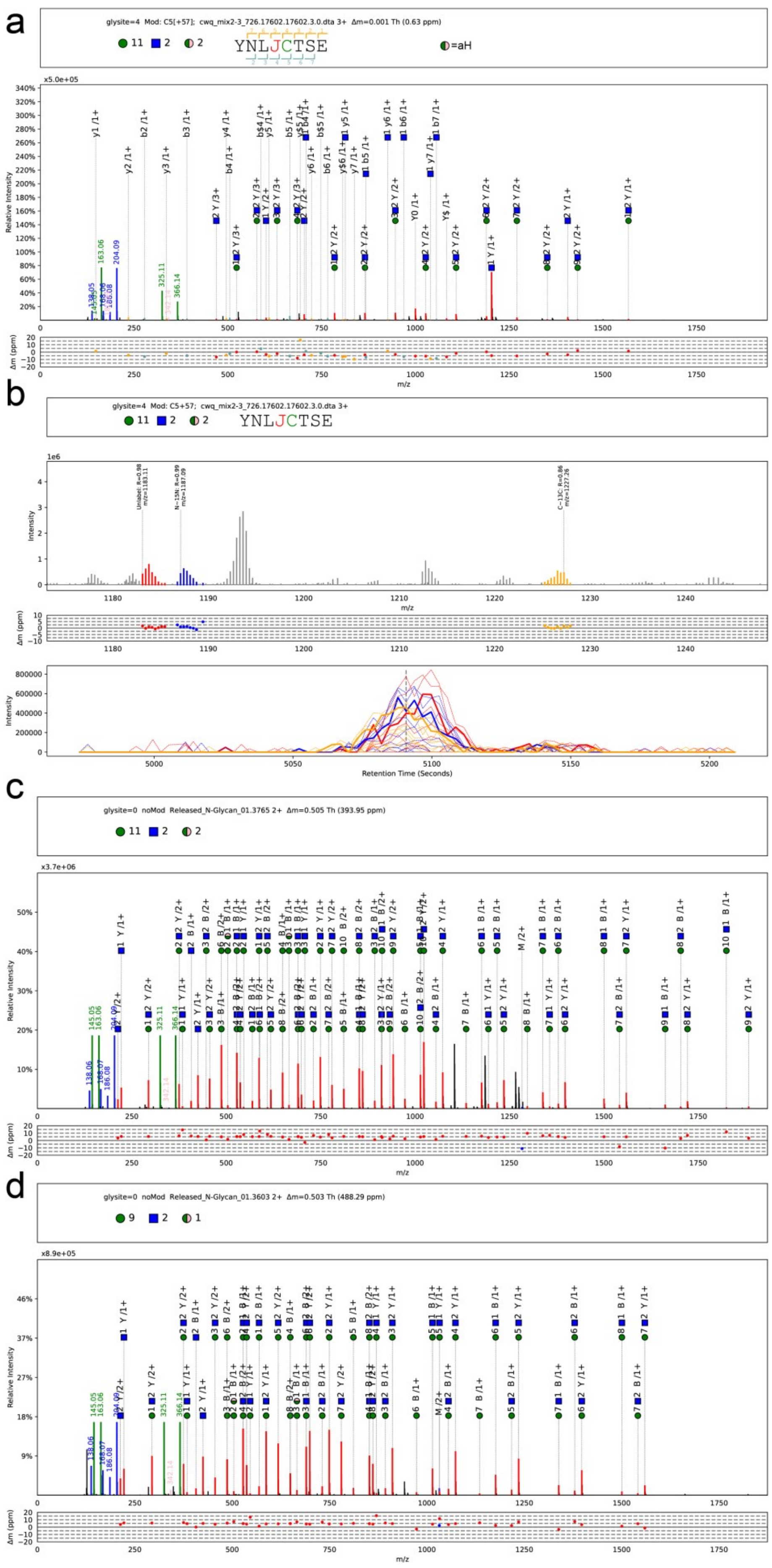

a
glysite=4 Mod: C5[+57]; cwq_mix2-3_726.17602.17602.3.0.dta 3+ Δm=0.001 Th (0.63 ppm)
11 2 2
YNLJCTSE
=aH
Relative Intensity
m/z
Δm (ppm)
b
glysite=4 Mod: C5+57; cwq_mix2-3_726.17602.17602.3.0.dta 3+
11 2 2
YNLJCTSE
Intensity
m/z
Δm (ppm)
Retention Time (Seconds)
c
glysite=0 noMod Released_N-Glycan_01.3765 2+ Δm=0.505 Th (393.95 ppm)
11 2 2
Relative Intensity
m/z
Δm (ppm)
d
glysite=0 noMod Released_N-Glycan_01.3603 2+ Δm=0.503 Th (488.29 ppm)
9 2 1
Relative Intensity
m/z
Δm (ppm)

**Supplementary Figure 4. sceHCD-MS/MS, $^{15}N/^{13}C$-labeled MS1, and EThcD MS/MS spectra of an O-Man glycopeptide with aH**

The O-Man glycopeptide is "SSSEAAASSSSESVAAASVEAER + Hex(8)aH(1)" in fission yeast samples. aH is represented by a half-greed-half-pink circle, meaning a modified Hex (green circle). (a) The sceHCD-MS/MS spectrum of this glycopeptide in $^{15}N/^{13}C$-labeled data. (b) The preceding MS1 spectrum of the sceHCD-MS/MS spectrum in (a). The unlabeled, $^{15}N$-labeled, and $^{13}C$-labeled precursors and their isotopes are annotated. All these isotope distributions are very similar to their theoretical isotope distributions. R is the calculated Pearson correlation coefficient between experimental and theoretical isotope distributions. The bottom figure in (b) is the extracted isotope patterns through retention time, wherein the colors for unlabeled, $^{15}N$-labeled, and $^{13}C$-labeled peaks are the same as they are at the top figure in (b). (c) EThcD-MS/MS spectrum of this O-Man glycopeptide in a sceHCD-pd-EThcD RAW file. The localized sites are: S1:H(1)@94, S2:H(1)@94, S3:H(1)@94, S8-S9:H(2)aH(1)@92, S10-S11:H(1)@90, S13:H(1)@90, S18:H(1)@90. @x means the SSGL probability is x%. These figures proved the existence of aH-O-Man glycopeptides in fission yeast samples.

(a) and (c) were directly and automatically annotated by gLabel (pGlyco3's built-in GPSM annotation tool) from the original MS/MS spectra (not deisotoped and deconvoluted). Annotation plots with multiple SSGL sites could be enabled in the pGlycoSite mode. (b) was generated by the in-house python script (gLabel-MS1).

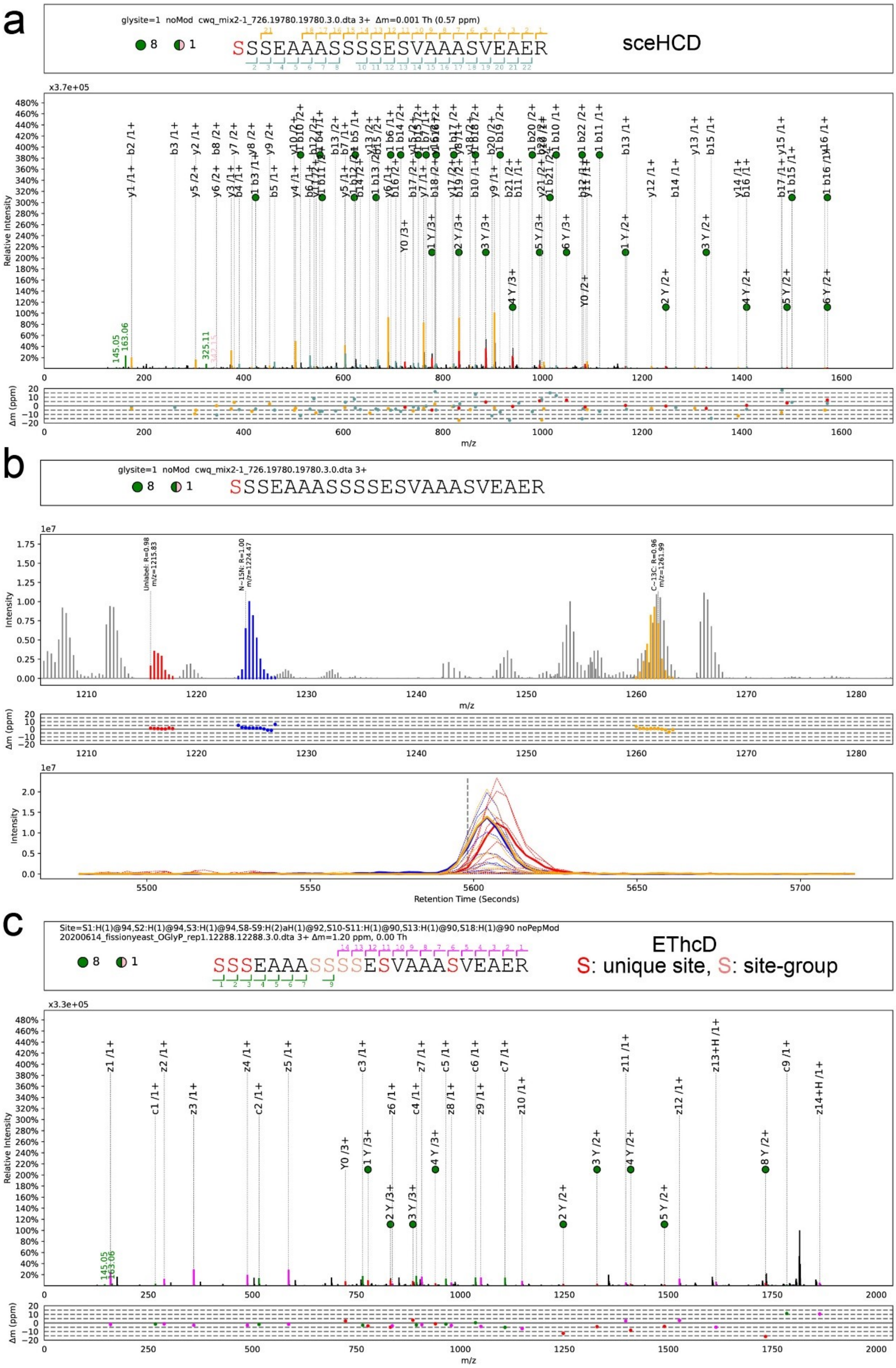

a
glysite=1 noMod cwq_mix2-1_726.19780.19780.3.0.dta 3+ Δm=0.001 Th (0.57 ppm)
8 1
SSSEAAASSSSESVAAASVEAER
sceHCD
x3.7e+05
Relative Intensity
Δm (ppm)
m/z
b
glysite=1 noMod cwq_mix2-1_726.19780.19780.3.0.dta 3+
8 1
SSSEAAASSSSESVAAASVEAER
Unlabel: R=0.98 m/z=1215.83
N−15N: R=1.00 m/z=1224.47
C−13C: R=0.96 m/z=1261.99
Intensity
m/z
Δm (ppm)
Retention Time (Seconds)
c
Site=S1:H(1)@94,S2:H(1)@94,S3:H(1)@94,S8-S9:H(2)aH(1)@92,S10-S11:H(1)@90,S13:H(1)@90,S18:H(1)@90 noPepMod
20200614_fissionyeast_OGlyP_rep1.12288.12288.3.0.dta 3+ Δm=1.20 ppm, 0.00 Th
8 1
SSSEAAASSSSESVAAASVEAER
EThcD
S: unique site, S: site-group
x3.3e+05
Relative Intensity
Δm (ppm)
m/z

**Supplementary Figure 5. N-glycosylation and O-Man analyses of the *ogm1* protein using all fission yeast datasets**

N-glycopeptides were analyzed using PXD005565 and PXD021128, while O-Man glycopeptides were analyzed using sceHCD-pd-EThcD data which were generated in this work.

The function of *ogm1* described in https://www.uniprot.org/uniprot/O13898 is: “transfers mannose from Dol-P-mannose to Ser or Thr residues on proteins. Required for normal cell growth and septum formation. Shown to actively O-mannosylate wsc1”. It would be quite confident that O-mannosylation occurs on this protein. We also found aH-contained N-glycosylation (site N443) and O-mannosylation. From this figure, we can see that aH-O-mannosylation may probably occur on the site S812 to S815 because there is no further evidence showing that aH occurs on other sites. H refers to Hex, and N refers to HexNAc.

*ogm1* (O13898)

| | | | | | | | | | | | | | | | | | | | | | | | | | |
|---|---|---|---|---|---|---|---|---|---|---|---|---|---|---|---|---|---|---|---|---|---|---|---|---|---|
| | | 443 | | | | 805 | 806 | 807 | | | | | 812 | 813 | 814 | 815 | | 817 | | | | | 822 | | |
| ... | S | N | G | ... | R | S | S | S | E | A | A | A | S | S | S | S | E | S | V | A | A | A | S | V | ... |

| Site | Glycan |
|---|---|
| N443 | H(9)N(2) |
| N443 | H(8)N(2) |
| N443 | H(10)N(2) |
| N443 | H(7)N(2) |
| N443 | H(7)N(2)aH(1) |
| N443 | H(8)N(2)aH(1) |
| N443 | H(8)N(2)F(1)aH(1) |
| N443 | H(6)N(2)F(1)aH(2) |

| Site | Glycan | Prob |
|---|---|---|
| S805 | H(1) | 0.94 |
| S806 | H(1) | 0.96 |
| S807 | H(1) | 0.96 |
| S805-S806 | H(2) | 0.93 |
| S806-S807 | H(2) | 0.91 |
| S805-S807 | H(3) | 0.92 |

| Site | Glycan | Prob |
|---|---|---|
| S812 | H(1) | 0.92 |
| S813 | H(1) | 0.92 |
| S814 | H(1) | 0.93 |
| S815 | H(1) | 0.93 |
| S815 | aH(1) | 0.91 |
| S812-S813 | H(3) | 0.90 |
| S812-S813 | H(2)aH(1) | 0.90 |
| S812-S814 | H(3) | 0.89 |
| S814-S815 | H(1) | 0.90 |
| S814-S815 | H(1)aH(1) | 0.90 |
| S813-S815 | H(2)aH(1) | 0.93 |
| S812-S815 | H(3)aH(1) | 0.93 |

| Site | Glycan | Prob |
|---|---|---|
| S817 | H(1) | 0.93 |
| S822 | H(1) | 0.96 |
| S812-S822 | H(5)aH(1) | 0.88 |
| S817-S822 | H(1) | 0.91 |
| S817-S817 | H(2) | 0.87 |

**Supplementary Figure 6. Comparison of pGlyco3 and pGlyco 2.0 on mouse-tissue datasets**

There are many differences by comparing pGlyco3 with pGlyco 2.0, including glycan database construction, glycan search (ion-indexing), glycan filtration, ETD supports, O-glycopeptide supports, pGlycoSite, and some fixed bugs. Some of them may cause different identification results. And the parameters of scoring schema were continuously fine-tuned using different datasets from different users in the development of pGlyco 2.0 (Liu et al. 2017[4]) to pGlyco3 during these years.

The scoring schema is: glycan score $Score_G = \sum_i \log(inten_i)\left(1-\left|\frac{merr_i}{tol_i}\right|^4\right) ratio_{ion}^{\alpha} ratio_{core}^{\beta}$, peptide score $Score_P = \sum_i \log(inten_i)\left(1-\left|\frac{merr_i}{tol_i}\right|^4\right) ratio_{ion}^{\gamma}$, and glycopeptide score $Score_{GP} = w \times Score_G + (1-w) \times Score_P$. In pGlyco 2.0, $\alpha = 0.56, \beta = 0.42, \gamma = 0.94, w = 0.35$. The final parameters in pGlyco3 were $\alpha = 0.56, \beta = 1.42, \gamma = 0.70, w = 0.35$.

It turns out that, these changes make the search better, as pGlyco3 could identify more N-glycopeptides in our previously published mouse-tissue datasets[4].

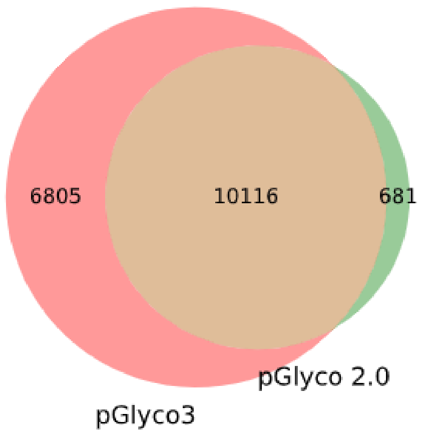

**Supplementary Figure 7. Software comparisons on HCD-pd-EThcD N-glycopeptide data**

We compared pGlyco3 with MSFragger and MetaMorpheus on HCD-pd-EThcD the N-glycopeptide dataset from MSV00083710[5]. MSV000083710 contains two datasets (a) human milk and (b) phospho-enriched Chinese hamster ovary cell (CHO). The venn diagrams were obtained based on unique glycopeptides. The search parameters are listed in Supplementary Data 1. pGlyco3 showed good identification on unique N-glycopeptides with sceHCD-pd-EThcD data. Furthermore, as pGlyco3 is specifically designed for the modified glycan search, it could identify more phospho-Hex (simplified as phoH) N-glycopeptides than the other two tools, as shown in figure b. All identified phoH-N-glycopeptides by pGlyco3 were supported by the phoH-diagnostic ion (243.026 m/z), hence they would be quite reliable. In (b), pGlyco3 can cover 48 of 49 phoH-identifications of MetaMorpheus, and one was lost due to the precursor detection error (c). among 275 uniquely identified phoH-glycopeptides of pGlyco3, 268 were because pGlyco3 could generated the phoH-glycans online, which were not in the search space of other tools. But the phoH-glycans of other 7 glycopeptides miss-identified by other tools were found in the 182-N-glycan database, an example of merged HCD+EThcD spectrum was shown in (d). There 7 miss-identifications may be also because of precursor detection errors of the other tools.

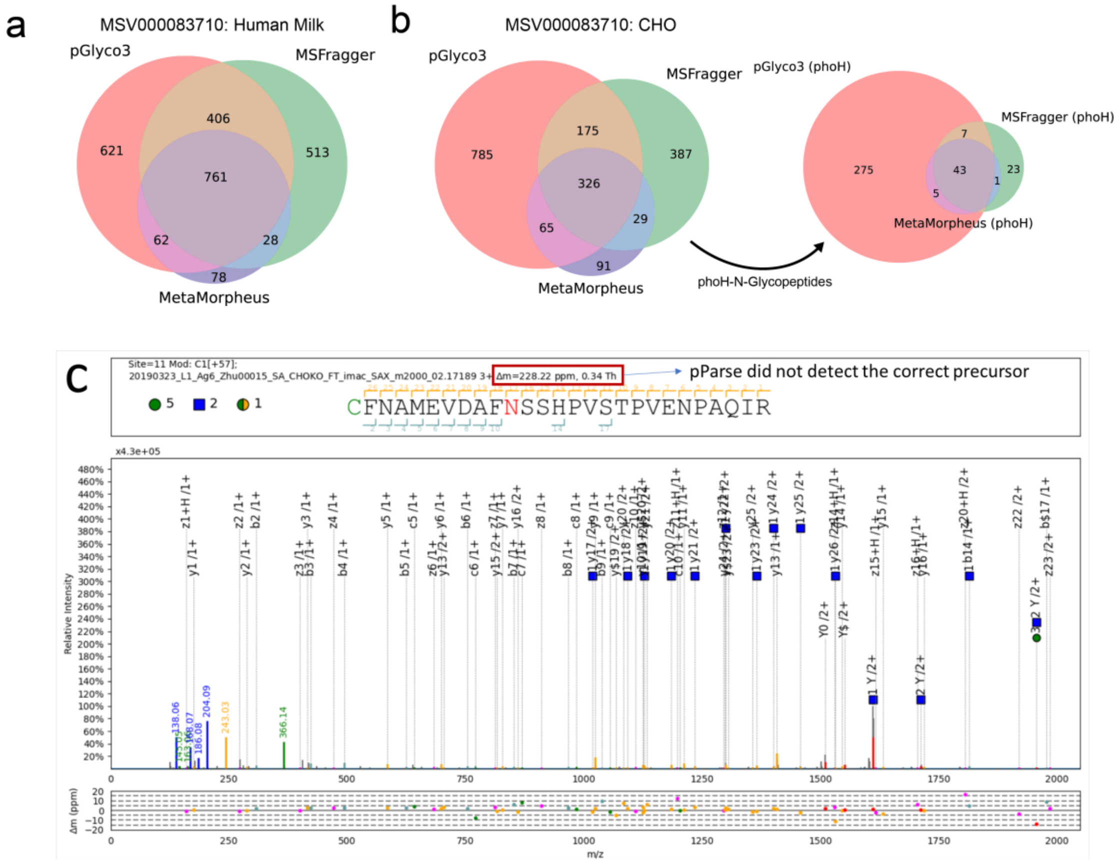

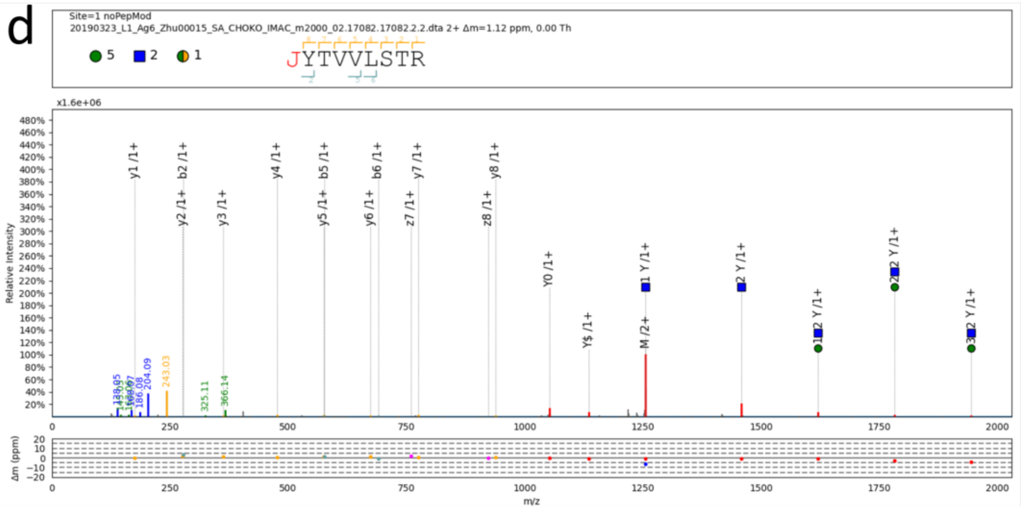
d
Site=1 noPepMod
20190323_L1_Ag6_Zhu00015_SA_CHOKO_IMAC_m2000_02.17082.17082.2.2.dta 2+ Δm=1.12 ppm, 0.00 Th
5
2
1
JYTVVLSTR
x1.6e+06
Relative Intensity
y1 /1+
b2 /1+
y2 /1+
y3 /1+
y4 /1+
b5 /1+
y5 /1+
b6 /1+
y6 /1+
y7 /1+
z7 /1+
y8 /1+
z8 /1+
Y0 /1+
Y$ /1+
M /2+
243.03
204.09
366.14
325.11
Δm (ppm)
m/z

**Supplementary Figure 8. aH in other public glycopeptide datasets**

aH-glycopeptides were also found in other public glycopeptide datasets with pGlyco3, showing the commonality of aH. Dataset information: Human serum N-glycopeptide dataset (InterVenn Serum, pGlyco-N-Human glycan database) is from ref[6]; AI-ETD mouse brain N-glycopeptide dataset (Riley AI-ETD, searched with pGlyco-N-Mouse glycan database) is from PXD011533[7]; N-glycopeptide datasets of standard proteins (Riley STD, pGlyco-N-Human glycan database) and HEK293 (Riley HEK293, pGlyco-N-Human glycan database) are from PXD017646[8]; EXoO datasets are from PXD009476[9], and searched with pGlyco-O-Glycan glycan database. A spectrum annotation example of an aH-glycopeptide from the EXoO-KidneyNormal dataset was show in (b).

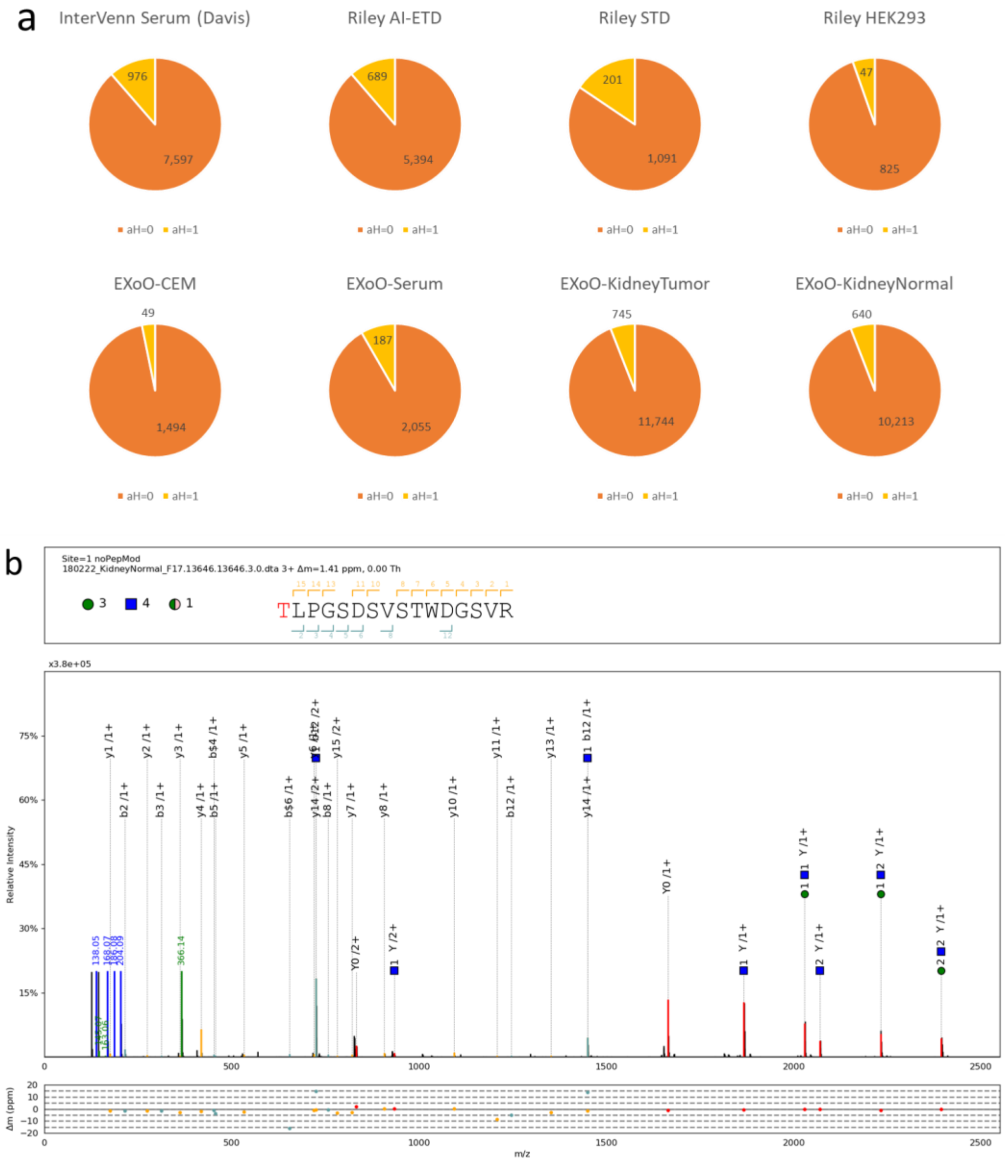

**Supplementary Figure 9. Search time and peptide FDR evaluation based on human serum O-glycopeptide data**

The Human serum O-glycopeptide data were searched by human protein database mixed with Arabidoposis (simplified as A), fussion yeast (simplified as Y) or worm (simplified as W). “Human+AYW” means pGlyco3 searches the protein database from human, A, Y, and W sequences. Even with a very large protein database (>10 million peptide sequences, 3 RAW files, searching MGF files parsed by pParse), pGlyco3 could finish the search and site localization with 43.8 minutes (subfigure a). And increasing the size of protein database would not much increase the peptide FDRs (subfigure b). All searches were tested using 3 processors with a Dell laptop with 32 GB physical memory.

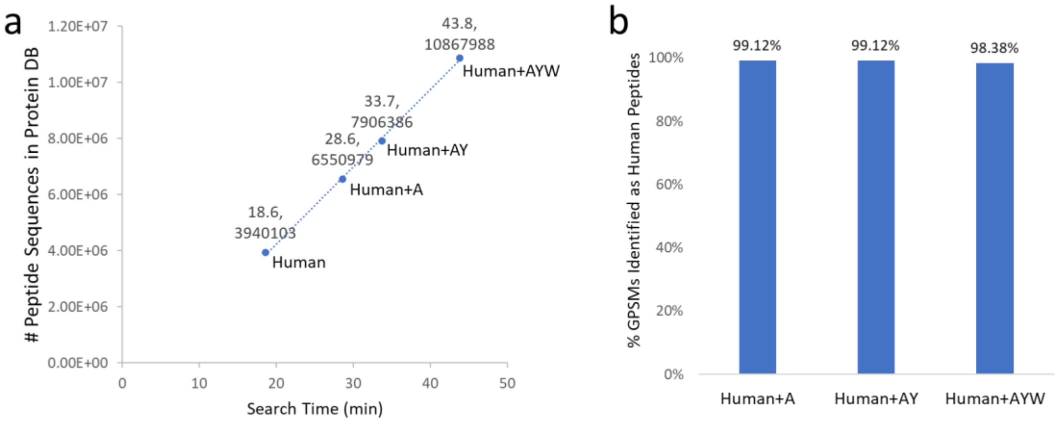

**Supplementary Figure 10. Spectrum annotation of the double-site N-glycopeptide “THTJ(H5N5A2F1)ISESHPJ(H5N2)ATF” of IgM, and its “Glu-C+trypsin” validation**

(a) EThcD (scan=3916) spectrum of “THTJ(H5N5A2F1)ISESHPJ(H5N2)ATF”; (b) HCD (scan=3915) spectrum of “THTJ(H5N5A2F1)ISESHPJ(H5N2)ATF”; (c) Spectrum of “Glu-C+trypsin” validation for “THTJ(H5N5A2F1)ISE”; (d) Spectrum of “Glu-C+trypsin” validation for “SHPJ(H5N2)ATFSAVGE”.

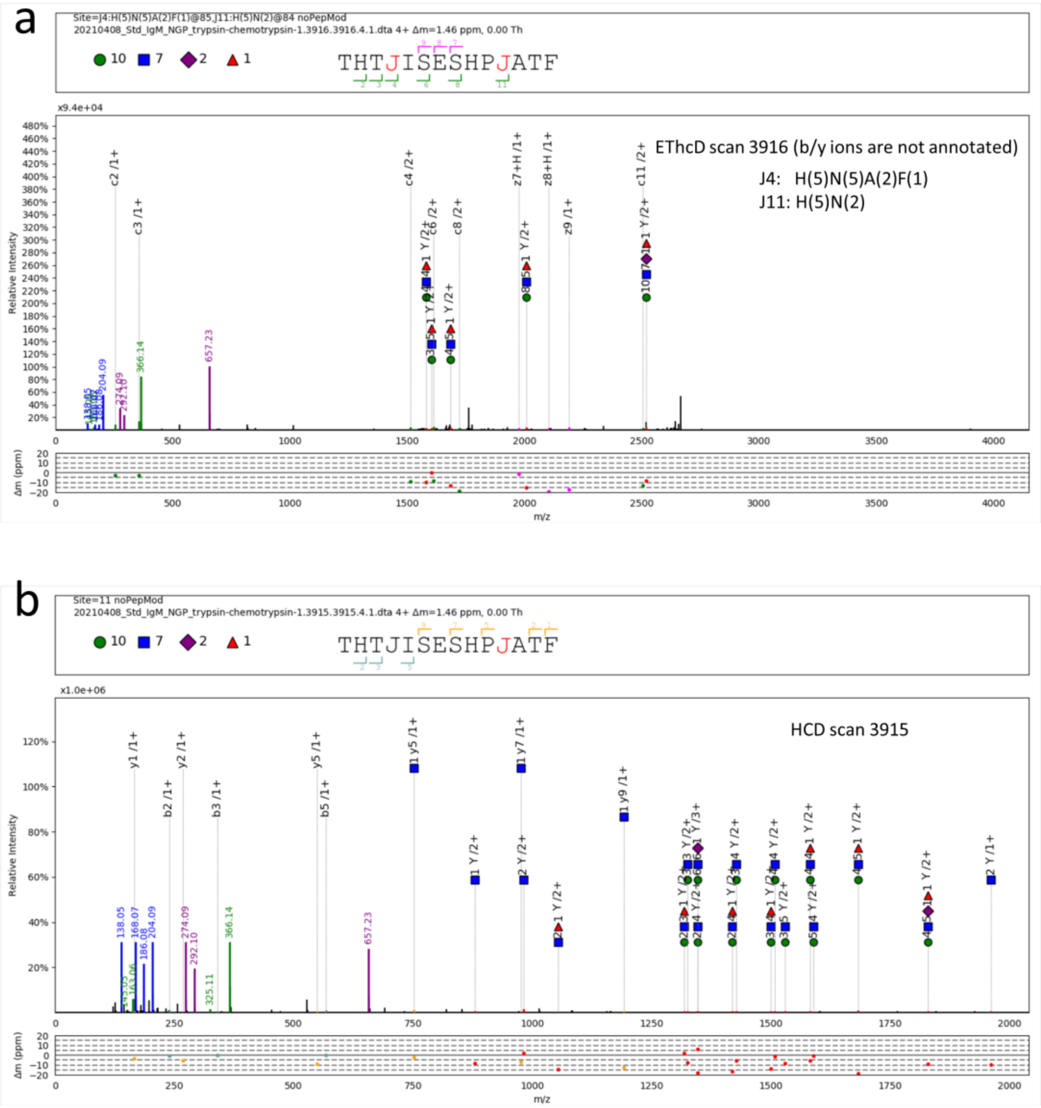

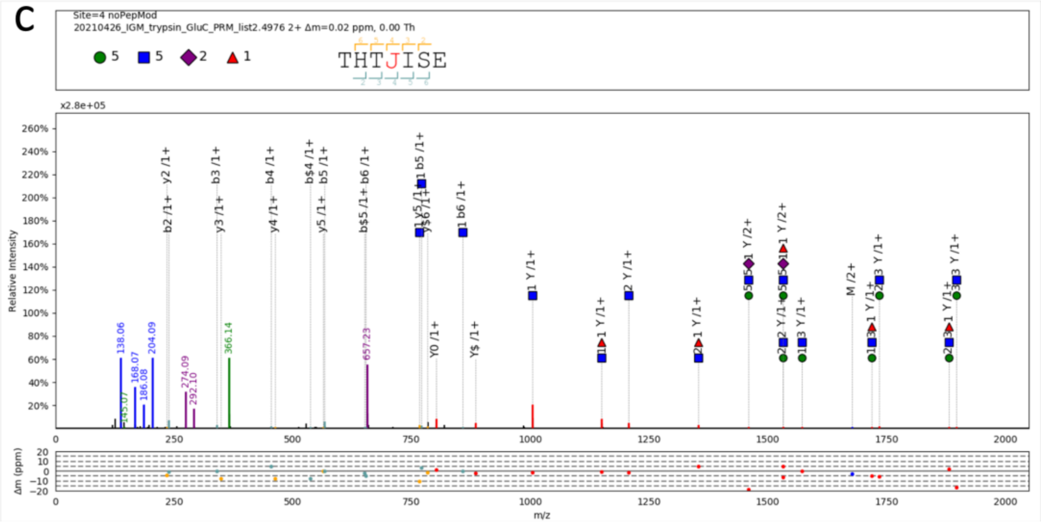
c
Site=4 noPepMod
20210426_IGM_trypsin_GluC_PRM_list2.4976 2+ Δm=0.02 ppm, 0.00 Th
THTJISE
x2.8e+05
Relative Intensity
Δm (ppm)
m/z

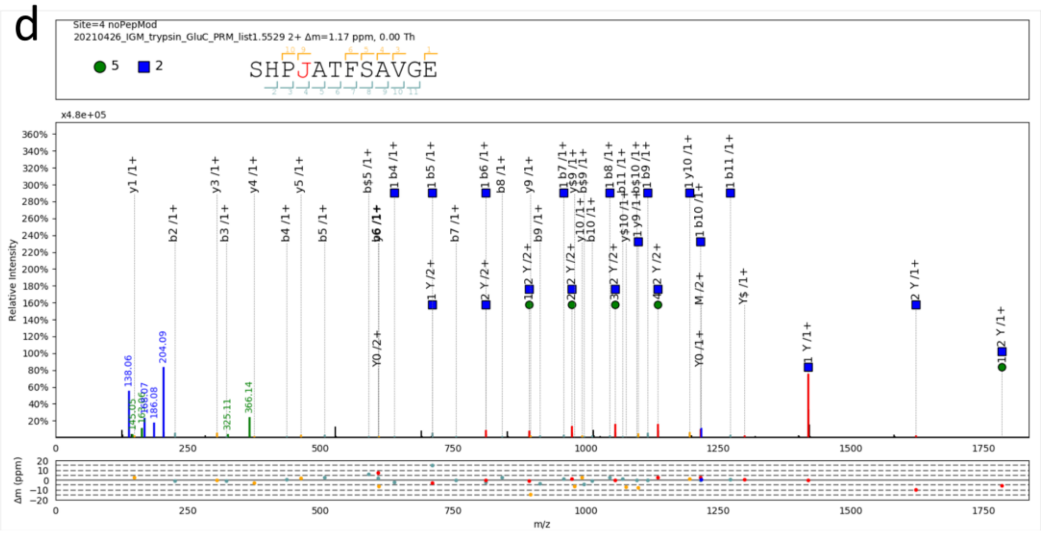
d
Site=4 noPepMod
20210426_IGM_trypsin_GluC_PRM_list1.5529 2+ Δm=1.17 ppm, 0.00 Th
SHPJATFSAVGE
x4.8e+05
Relative Intensity
Δm (ppm)
m/z

**Supplementary Figure 11. HCD spectrum of the example in Fig. 3f**

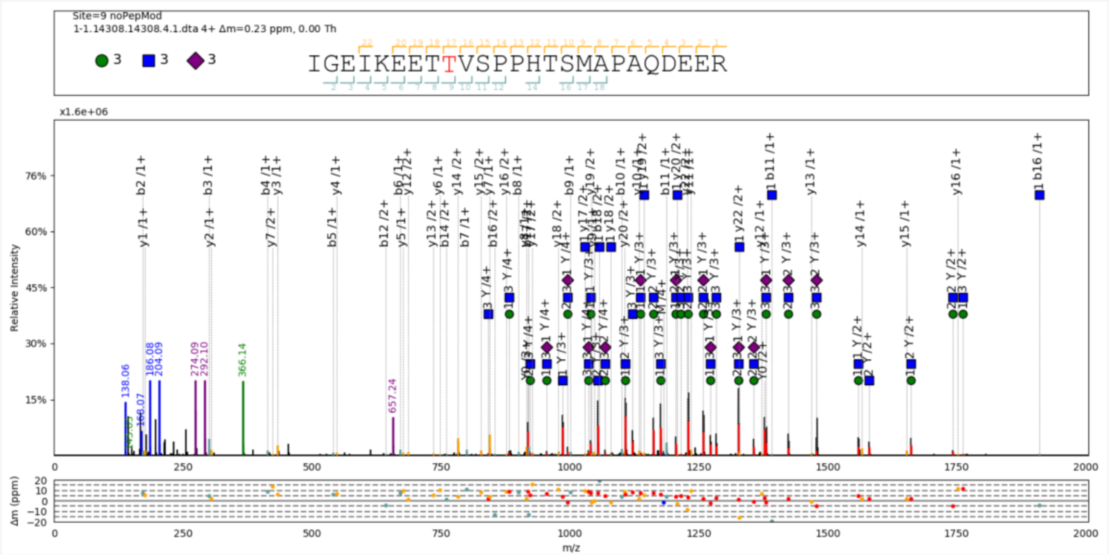

### Supplementary Note 1. Canonicalization-based modified glycan generation

First of all, we listed pGlyco3's built-in glycan databases in Supplementary Note Table 1.

Supplementary Note Table 1. Glycan database sources

| GDB Name | # Glycan Compositions (Structures) | Source |
|---|---|---|
| pGlyco-N-Human | 695 (2921) | Combination of N-glycans and their sub-structures in Glycome-DB[10] with the sub-structures of three largest high-mannose, hybrid, and complex N-glycans. See ref[11]. |
| pGlyco-N-Human-multi | 2051 (4278) | Combination of 1 to 2 pGlyco-N-Human databases, for multi-site N-glycopeptides. Redundancies were removed |
| pGlyco-N-Mouse-Large | 1622 (7878) | Extending NeuAc to NeuGc in pGlyco-N-Human |
| pGlyco-N-Mouse | 1234 (6662) | Glycans with size > 20 were removed from pGlyco-N-Mouse-Large |
| pGlyco-N-Plant | 107 (289) | N-glycans and their sub-structures of three largest high-mannose, hybrid, and complex N-glycans in plants. See https://en.wikipedia.org/wiki/N-linked_glycosylation. |
| pGlyco-O-Glycan | 156 (552) | O-GalNAc glycans and their sub-structures displayed in published works[12, 13] and wikipedia (https://en.wikipedia.org/wiki/O-linked_glycosylation). |
| Multi-Site-O-Glycan | 445 (6177) | Combination of 1 to 8 O-GalNAc glycans displayed in ref[12], the combined glycan sizes were limited to 20. Redundancies were removed. pGlycoSite is not limited by 8 O-glycosites thanks to the dynamic |

| | | |
|---|---|---|
| | | programming algorithm. |

All these glycan databases were compiled into canonicalization-based databases in pGlyco3 (.gdb files). Tree canonicalization is not a new algorithm, it has been frequently used in computer science to mine tree-structure data[14], and GlycoWorkbench also encodes glycans as canonicalization. The main reason for pGlyco3 to use canonicalization to encode glycan structures is that it is easy to generate modified glycans based on canonical strings, as shown in Supplementary Note Figure 1.

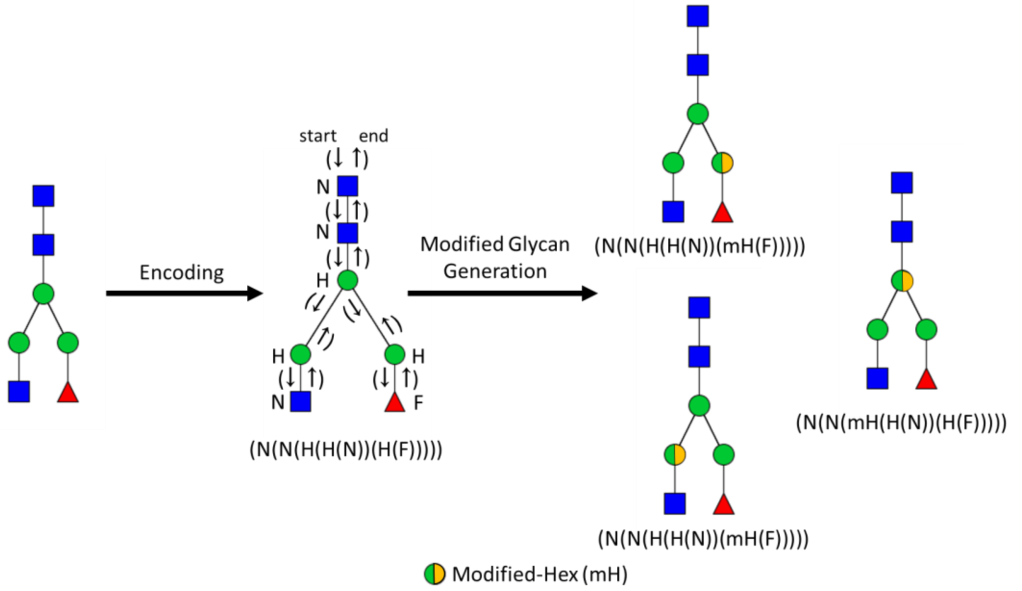

Supplementary Note Figure 1. An example of canonicalization-based glycan structure encoding and modified glycan generation. Modified glycans could be generated by replacing H with mH in the encoded strings.

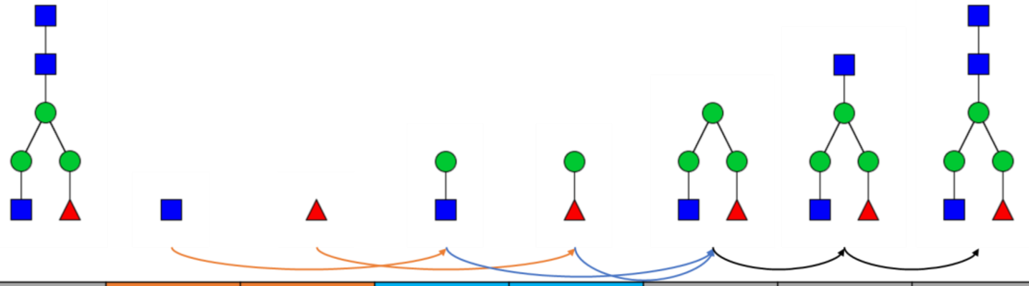

| DFS step | Step 1 | Step 1 | Step 2 | Step 2 | Step 3 | Step 4 | Step 5 |
|---|---|---|---|---|---|---|---|
| Y Fragments of subtrees | N(1) | F(1) | H(1)<br>H(1)N(1) | H(1)<br>H(1)F(1) | H(1)<br>H(2)<br>H(3)<br>H(2)N(1)<br>H(2)F(1)<br>H(3)N(1)<br>H(3)F(1)<br>H(3)N(1)F(1) | N(1)<br>H(1)N(1)<br>H(2)N(1)<br>H(3)N(1)<br>H(2)N(2)<br>H(2)N(1)F(1)<br>H(3)N(2)<br>H(3)N(1)F(1)<br>H(3)N(2)F(1) | N(1)<br>N(2)<br>H(1)N(2)<br>H(2)N(2)<br>H(3)N(2)<br>H(2)N(3)<br>H(2)N(2)F(1)<br>H(3)N(3)<br>H(3)N(2)F(1)<br>H(3)N(3)F(1) |

Supplementary Note Figure 2. An example of the DFS algorithm to generate Y ions for a given glycan tree. Y ions from two subtrees in step 2 are joined into their parent node to generate Y ions in step 3.

The encoding algorithm is quite simple, as illustrated in Supplementary Algorithm 1. The decoding algorithm is similar. pGlyco3 also uses the depth-first search (DFS) algorithm to generate corresponding Y ions of the given glycan (Supplementary Algorithm 1 and Supplementary Note Figure 2).

As there are more than one glycan trees for multi-site N- and O-glycans, all Y ions including the parent glycan composition from different trees are cross-joined and redundant Y ions are removed (similar to Supplementary Note Figure 2). For example, for two glycans, say (N(H(A))) and (N(H)), Y ions of (N(H(A))) are Y0, N(1), N(1)H(1), and N(1)H(1)A(1); Y ions of (N(H)) are Y0, N(1), and N(1)H(1). After cross-joining, the non-redundant Y ions would be Y0, N(1), N(2), N(1)H(1), N(2)H(1), N(2)H(2), N(1)H(1)A(1), N(2)H(1)A(1), and N(2)H(2)A(1), where N(2)H(2)A(1) is the parent composition of the joined glycan.

Supplementary Algorithm 1. Pseudo code of canonicalization-based glycan encoding algorithm.

DFS: Depth-First Search;

glycan_tree: the tree structure of a given glycan

```
DFS_Glycan_Encoding(glycan_tree):
    subtree_codes = list()
    root = Get_Root_Glyco(glycan_tree)
    foreach subtree of glycan_tree:
        subtree_codes.append(DFS_Canon_Encoding(subtree))
    Sort subtree_codes
    concat_subtree_codes = String_Concatenate(subtree_codes)
    codes = "(" + root + concat_subtree_codes + ")"
    Return codes
```

In pGlyco3, glycan structures are essential to generate possible glycan Y ions and determine correct core Y ions for glycan search to reduce the possibility of mismatches, but it does not mean that we can identify the glycan structure from the MS/MS spectrum of an intact glycopeptide. There are some reasons for the difficulties of structure identification from the glycopeptide MS/MS spectrum: 1) Many glycan structures share the identical Y ions or only differ in few Y ions, for example, the glycan "(N(N(H(H))(H(H(H)))))" and "(N(N(H)(H(H(H(H))))))"; 2) Because glycans and peptides are large, the masses of some Y ions (attached with peptides) are too large to detect in the MS/MS spectra, especially from the Orbitrap-based instruments; 3) Glycan rearrangement in MS/MS[15]. Therefore, pGlyco3 reports the glycan compositions as well as the "plausible structures" for the identified glycopeptides. The presence of oxonium ions may also provide some information to determine the structure.

### Supplementary Note 2. Glycan ion-indexing

Glycan ion-indexing is designed for the fast glycan scoring and filtration for the glycan-first search. Note that glycans are searched before peptides in the glycan-first search workflow, hence we cannot directly calculate the Y ions (glycan Y ions plus the peptide mass) of a glycopeptide for glycan scoring. But, for a deconvoluted spectrum, if a peak is a Y ion, then "precursor mass – peak mass" would be the mass complementary to the Y ion which does not contain the peptide parts while keeps the glycan fragment information. The key observation is:

$$peak\ mass = peptide\ mass + glycan\ Y\ ion\ mass + peak\ mass\ error(+proton)$$

$$precursor\ mass = peptide\ mass + glycan\ mass + precursor\ mass\ error(+proton)$$

$$\begin{aligned} \Rightarrow precursor\ mass - peak\ mass &= glycan\ mass - glycan\ Y\ ion\ mass + \ mass\ error \\ &= Y\text{-}complementary\ mass + mass\ error \end{aligned}.$$

Based on this observation, we designed a glycan ion-indexing algorithm for the fast glycan-first search. An example of the glycan ion-indexing structure is shown in Supplementary Note Figure 3. Glycan ion-indexing is built as follows:

1. Load glycan structures and generate modified glycans;
2. Generate all Y ions for each structure, and also checks if the Y ion is the core ion (Supplementary Table 2). The Y0 ion is also considered as the core Y ion, as Y0 ion represents the peptide mass in a glycopeptide;
3. Generate corresponding Y-complementary ions of Y ions for each glycan structure;
4. Merge all Y-complementary ions for all glycans, and build an indexing table for Y-complementary ions and their originating glycan IDs. pGlyco3 uses an extra bit in the 32-bit integer of the glycan ID to record whether the Y-complementary ion is from core Y ions or not. The bit-level operation could not only save the memory but also enable us to simultaneously score the Y ion and its core Y ion for the glycan(s) once a Y-complementary is matched;
5. Sort the Y-complementary ion-indexing table by the Y-complementary ion masses. Build a hash table for the Y-complementary ion-indexing table (0.01 Da per bin in the hash table) to enable O(1) query time.

**The time complexity for searching an MS/MS spectrum via the Y-complementary ion-indexing table is O(#peaks).** Initialize all glycan scores and the core scores to zero. For a given peak in an MS/MS spectrum, calculate the query mass as "precursor mass – peak mass", and then search the query mass via the Y-complementary ion-indexing table, if a Y-complementary ion is matched, add the ion score (counting score) to its originating glycan IDs, and also add the core ion score to its originating glycan IDs if the core-bit is 1. Querying a peak only takes O(1) time by using the glycan ion-indexing, hence the time complexity for querying all peaks of the spectrum is O(# peaks).

1. Core: Is the Y ion the trimannosyl-core ion of this glycan? Y=yes, N=no
2. Y-complementary ion = glycan composition – Y ion

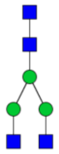

Glycan ID = i

| Glycan Y Ion | Core? | Y-Complementary Ion |
|---|---|---|
| Y0 | Y | HexNAc(4)Hex(3) |
| HexNAc(1) | Y | HexNAc(3)Hex(3) |
| HexNAc(2) | Y | HexNAc(2)Hex(3) |
| HexNAc(2)Hex(1) | Y | HexNAc(2)Hex(2) |
| HexNAc(2)Hex(2) | Y | HexNAc(2)Hex(1) |
| HexNAc(2)Hex(3) | Y | HexNAc(2) |
| HexNAc(3)Hex(2) | N | HexNAc(1)Hex(1) |
| HexNAc(3)Hex(3) | N | HexNAc(1) |

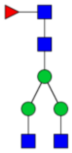

Glycan ID = j

| Glycan Y Ion | Core? | Y-Complementary Ion |
|---|---|---|
| Y0 | Y | HexNAc(4)Hex(3)Fuc(1) |
| HexNAc(1) | Y | HexNAc(3)Hex(3)Fuc(1) |
| HexNAc(2) | Y | HexNAc(2)Hex(3)Fuc(1) |
| HexNAc(2)Hex(1) | Y | HexNAc(2)Hex(2)Fuc(1) |
| HexNAc(2)Hex(2) | Y | HexNAc(2)Hex(1)Fuc(1) |
| HexNAc(2)Hex(3) | Y | HexNAc(2)Fuc(1) |
| HexNAc(3)Hex(2) | N | HexNAc(1)Hex(1)Fuc(1) |
| HexNAc(3)Hex(3) | N | HexNAc(1)Fuc(1) |
| HexNAc(4)Hex(3) | N | Fuc(1) |
| HexNAc(1)Fuc(1) | Y | HexNAc(3)Hex(3) |
| HexNAc(2)Fuc(1) | Y | HexNAc(2)Hex(3) |
| HexNAc(2)Hex(1)Fuc(1) | N | HexNAc(2)Hex(2) |
| HexNAc(2)Hex(2)Fuc(1) | N | HexNAc(2)Hex(1) |
| HexNAc(2)Hex(3)Fuc(1) | N | HexNAc(2) |
| HexNAc(3)Hex(2)Fuc(1) | N | HexNAc(1)Hex(1) |
| HexNAc(3)Hex(3)Fuc(1) | N | HexNAc(1) |

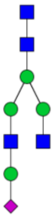

Glycan ID = k

| Glycan Y Ion | Core? | Y-Complementary Ion |
|---|---|---|
| Y0 | Y | HexNAc(4)Hex(4)NeuAc(1) |
| HexNAc(1) | Y | HexNAc(3)Hex(4)NeuAc(1) |
| HexNAc(2) | Y | HexNAc(2)Hex(4)NeuAc(1) |
| HexNAc(2)Hex(1) | Y | HexNAc(2)Hex(3)NeuAc(1) |
| HexNAc(2)Hex(2) | Y | HexNAc(2)Hex(2)NeuAc(1) |
| HexNAc(2)Hex(3) | Y | HexNAc(2)Hex(1)NeuAc(1) |
| HexNAc(3)Hex(2) | N | HexNAc(1)Hex(2)NeuAc(1) |
| HexNAc(3)Hex(3) | N | HexNAc(1)Hex(1)NeuAc(1) |
| HexNAc(4)Hex(3) | N | Hex(1)NeuAc(1) |
| HexNAc(4)Hex(4) | N | NeuAc(1) |
| HexNAc(3)Hex(3)NeuAc(1) | N | HexNAc(1)Hex(1) |
| HexNAc(3)Hex(4)NeuAc(1) | N | HexNAc(1) |

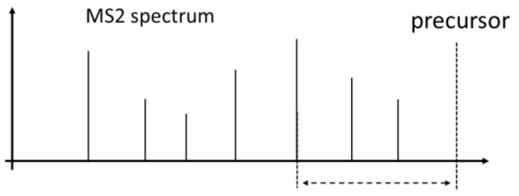

**query mass** =
precursor mass – peak mass

Searching **query mass** against Y-complementary ions of all glycans within O(1) time

**Hashed Glycan Ion-Indexing Table**

| Y-Complementary Ions | Mass | Glycan ID&Core |
|---|---|---|
| Fuc(1) | 146.058 | j,N; |
| HexNAc(1) | 203.079 | i,N; j,N; k,N; |
| NeuAc(1) | 291.095 | k,N; |
| HexNAc(1)Fuc(1) | 349.137 | j,N; |
| HexNAc(1)Hex(1) | 365.131 | i,N; j,N; k,N; |
| HexNAc(2) | 406.158 | i,Y; j,N; |
| Hex(1)NeuAc(1) | 453.147 | k,N; |
| HexNAc(1)Hex(1)Fuc(1) | 511.189 | j,N; |
| HexNAc(2)Fuc(1) | 552.216 | j,Y; |
| HexNAc(2)Hex(1) | 568.21 | i,Y; j,N; |
| HexNAc(1)Hex(1)NeuAc(1) | 656.226 | k,N; |
| HexNAc(2)Hex(1)Fuc(1) | 714.268 | j,Y; |
| HexNAc(2)Hex(2) | 730.262 | i,Y; j,N; |
| HexNAc(1)Hex(2)NeuAc(1) | 818.278 | k,N; |
| HexNAc(2)Hex(1)NeuAc(1) | 859.305 | k,Y; |
| HexNAc(2)Hex(2)Fuc(1) | 876.32 | j,Y; |
| HexNAc(2)Hex(3) | 892.314 | i,Y; j,Y; |
| HexNAc(2)Hex(2)NeuAc(1) | 1021.357 | k,Y; |
| HexNAc(2)Hex(3)Fuc(1) | 1038.372 | j,Y; |
| HexNAc(3)Hex(3) | 1095.393 | i,Y; j,Y; |
| HexNAc(2)Hex(3)NeuAc(1) | 1183.409 | k,Y; |
| HexNAc(3)Hex(3)Fuc(1) | 1241.451 | j,Y; |
| HexNAc(4)Hex(3) | 1298.472 | i,Y; |
| HexNAc(2)Hex(4)NeuAc(1) | 1345.461 | k,Y; |
| HexNAc(4)Hex(3)Fuc(1) | 1444.53 | j,Y; |
| HexNAc(3)Hex(4)NeuAc(1) | 1548.54 | k,Y; |
| HexNAc(4)Hex(4)NeuAc(1) | 1751.619 | k,Y; |

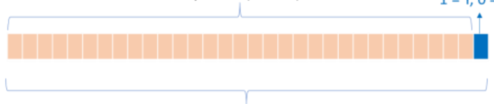

Supplementary Note Figure 3. An example of glycan ion-indexing structure.

### Supplementary Note 3. The pGlycoSite algorithm

#### The time complexity of the brute-force SSGL method

With Y ions and b/y/c/z ions, it is not difficult to identify the peptide part and the total glycan composition for a given HCD-pd-ETxxD spectrum. Hence the major problem is, for a given peptide and glycan composition, how many glycopeptide-forms (isoforms) are there in total? To make this question simpler, we assume the glycan composition only contains one monosaccharide-type (for example, only Hex for O-Man Glycan). Suppose that there are $S$ candidate glycosylation sites on the peptide sequence and $|G|$ monosaccharides in the glycan composition. The central problem is to calculate how many different ways to put $|G|$ balls (monosaccharides) into $S$ bins (sites). Note that each bin (site) may contains $0, 1, \dots, |G|$ balls (monosaccharides). This problem could be solved by the stars-and-bars method (https://en.wikipedia.org/wiki/Stars_and_bars_(combinatorics)). Hence the total isoforms will be $C_{S+|G|-1}^{S-1}$, which is exponentially increased as $S$ or $|G|$ increases, as shown in Supplementary Note Table 2. For each isoform, we have to calculate its c/z ions to match against the spectrum, hence the time complexity is $O\left((L-1) \times C_{S+|G|-1}^{S-1}\right)$, which could be simplified as $O\left(L \times C_{S+|G|-1}^{S-1}\right)$, where $L$ is the peptide length. If there are $T$ monosaccharide types, the number of isoforms will be $\prod_{i=1}^{T} C_{S+G_i-1}^{G_i-1}$ in the worst case, where $G_i$ is the monosaccharide number of the $ith$ monosaccharide type, hence the time complexity upper bound would be $O(L \times \prod_{i=1}^{T} C_{S+G_i-1}^{G_i-1})$.

Therefore, for a given peptide and glycan composition, if we use brute-force method to localize the sites using the ETxxD spectrum, the time complexity is $O(L \times \prod_{i=1}^{T} C_{S+G_i-1}^{G_i-1})$. It is feasible for the small glycans with not too many candidate sites, but it would be very slow for large glycans with dense (e.g., mucin-type) sites.

Supplementary Note Table 2. The values of $C_{S+G-1}^{S-1}$ with different $S$ and $G$.

| $S$ | $G$ | # isoforms | $S$ | $G$ | # isoforms |
|---|---|---|---|---|---|
| 7 | 11 | 12,376 | 11 | 7 | 19,448 |
| 8 | 11 | 31,824 | 11 | 8 | 43,758 |
| 9 | 11 | 75,582 | 11 | 9 | 92,378 |
| 11 | 11 | 352,716 | 11 | 11 | 352,716 |
| 13 | 11 | 1,352,078 | 11 | 13 | 1,144,066 |
| 16 | 11 | 7,726,160 | 11 | 16 | 5,311,735 |

The pGlycoSite algorithm is designed to avoid the brute-force enumeration. Before introducing pGlycoSite, we firstly define the "sub-glycan". In pGlyco3, each glycan is represented by a vector, for example, if there are $T$ monosaccharide types, and the number of the $ith$ monosaccharide type is $G_i$, then glycan $G$ could be represented as a vector $(G_1, G_2, \dots, G_T)$. The glycan $g = (g_1, g_2, \dots, g_T)$ is the sub-glycan of $G$ if $g_i \le G_i\ \forall i = 1, \dots, T$. For a given glycan $G = (G_1, G_2, \dots, G_T)$, $F$ is defined as the number of sub-glycan compositions (forms) of $G$, which is: $F = \prod_{i=1}^{T}(G_i + 1)$.

The key observation for pGlycoSite algorithm is that, no matter how many isoforms there are for a given peptide and glycan composition, there are at most $F \times (L-1)$ c and z ions, this is because different glycopeptide-isoforms probably share a large proportion of same c/z ions. We just need to generate all possible c/z ions instead of all possible isoforms, as shown in Supplementary Note Figure 4a. These c/z ions are then matched against the ETxxD spectrum, as shown in Supplementary Note Figure 4b ($ScoreTable$). If both c/z ions in a cell are not matched, pGlycoSite will further consider c-H/z+H ions. pGlycoSite uses a dynamic programming algorithm to obtain the best score from bottom left to up right (Supplementary Note Figure 4c):

$$BestPath[g,p] = \begin{cases} \max\limits_{\forall g_s \le g} BestPath[g_s, p-1] + ScoreTable\,[g,p] & if\ IsValidPath(g_s, g, p) \\ \times & if\ not\ \exists g_s \le g\ \ IsValidPath(g_s, g, p) \end{cases},$$

where $g$ refers to a sub-glycan composition of $G$, $p$ refers to $p$th position of the peptide sequence. $g_s < g$ means that $g_s$ is the sub-glycan of $g$. $IsValidPath(g_s, g, p)$ is designed to check whether the path from $[g_s, p-1]$ to $[g,p]$ is valid or not. Assume $G$ is the identified glycan composition, then a path from $[g_s, p-1]$ to $[g,p]$ is **invalid** if one criterion of follows is met:

1. not $g_s \le g$;
2. $g_s < g$ but $p-1$ is not a candidate glycosylation site;
3. $g - g_s$ or $G - g$ is not a legal glycan combination;
4. $g < G$, but there are no candidate sites after site $p-1$.

Otherwise, the path from $[g_s, p-1]$ to $[g,p]$ is valid. There are two different ways to do step 3: a) check if $g - g_s$ and $G - g$ are in the glycan database; b) check if $g - g_s$ and $G - g$ contain the reducing-end monosaccharide (HexNAc for N- and O-glycosylation, and Hex for O-mannosylation). The method (a) is recommended as it can avoid many non-sense paths, and hence will be more accurate in principle. It could be enabled by "pGlycoSite: Localized Glycans Must Be in GDB" check-box in pGlyco3's GUI. We used the method (a) by default in this manuscript. For the pGlycoSite validation in Fig. 3d and Supplementary Fig. 2, we used the method (b). unless specially mentioned. We pre-build a glycan combination table of the glycan database before localization, enabling $O(1)$ checking time. After dynamic programming, $BestPath[G, L]$ then stores the best path score. pGlycoSite backtracks the $BestPath$ table to obtain the best path(s). Site $p-1$ contains a glycan $g - g_s$ if $g_s < g$. If there are multiple paths reaching the same best path score, sites from the branch position to the merging position are regarded as a “site-group”, as shown in Supplementary Note Figure 4d (S3~T5). A site-group means that we could not further distinguish sites in the group by using the matched c/z ions in the current scoring schema.

**The time complexity of the pGlycoSite algorithm is** $O(L \times F^2)$. From the dynamic programming equation, we can see that, 1) calculating $BestPath[g,p]$ only needs to check the $(p-1)$th amino acid, hence the time complexity is $O(L)$ in the peptide dimension; 2) calculating $BestPath[g,p]$ needs to check all $g_s$, and we have to visit all $g$, hence the time complexity is $O(F^2)$ in the glycan dimension. The total time complexity is $O(L \times F^2)$. The time complexities of the brute-force method and pGlycoSite are listed in Supplementary Note Table 3.

Supplementary Note Table 3. Time complexity comparisons.

| $S$ | $\lvert G \rvert$ | Brute-force complexity | pGlycoSite complexity |
|---|---|---|---|
| 11 | 7 | 19,448× $L$ | 49× $L$ |
| 11 | 8 | 43,758× $L$ | 64× $L$ |
| 11 | 9 | 92,378× $L$ | 81× $L$ |
| 11 | 11 | 352,716× $L$ | 121× $L$ |
| 11 | 13 | 1,144,066× $L$ | 169× $L$ |
| 11 | 16 | 5,311,735× $L$ | 256× $L$ |

**a**

Enumerated c/z Ion Table

| (H, N) | | | | | | | | | | | | | |
|---|---|---|---|---|---|---|---|---|---|---|---|---|---|
| (2, 2) | c/z | c/z | c/z | c/z | c/z | c/z | c/z | c/z | c/z | c/z | c/z | c/z | c/z |
| (2, 1) | c/z | c/z | c/z | c/z | c/z | c/z | c/z | c/z | c/z | c/z | c/z | c/z | c/z |
| (2, 0) | c/z | c/z | c/z | c/z | c/z | c/z | c/z | c/z | c/z | c/z | c/z | c/z | c/z |
| (1, 2) | c/z | c/z | c/z | c/z | c/z | c/z | c/z | c/z | c/z | c/z | c/z | c/z | c/z |
| (1, 1) | c/z | c/z | c/z | c/z | c/z | c/z | c/z | c/z | c/z | c/z | c/z | c/z | c/z |
| (1, 0) | c/z | c/z | c/z | c/z | c/z | c/z | c/z | c/z | c/z | c/z | c/z | c/z | c/z |
| (0, 2) | c/z | c/z | c/z | c/z | c/z | c/z | c/z | c/z | c/z | c/z | c/z | c/z | c/z |
| (0, 1) | c/z | c/z | c/z | c/z | c/z | c/z | c/z | c/z | c/z | c/z | c/z | c/z | c/z |
| (0, 0) | c/z | c/z | c/z | c/z | c/z | c/z | c/z | c/z | c/z | c/z | c/z | c/z | c/z |

T P S P T V A H E S N W A K
(5 under T5, 10 under S10)

Table interpretation

Peptide: TPSPTVAHESNWAK
Glycan: H(2)N(2), H=Hex, N=HexNAc
Each row represents a sub-glycan, each column represents a peptide's cleavage position. The c ion of each cell refers to N-term residues of its column plus the sub-glycan of its row. The z ion is the complementary of c ion.

In this cell:
c ion = TPSPTVAHESN (c11) + H(1)
z ion = WAK (z3) + H(1)N(2)

Matching against the spectrum (Fig. 3a)

**b**

Matched Enumerated c/z Ion Table (ScoreTable)

| (H, N) | | | | | | | | | | | | |
|---|---|---|---|---|---|---|---|---|---|---|---|---|
| (2, 2) | | | | z | c&z | c&z | c&z | c&z | c&z | c&z | c&z | z |
| (2, 1) | | | | z | c | c | | | | | | |
| (2, 0) | | | | | | | | | | | | |
| (1, 2) | | z | | | c | | | | z | | | |
| (1, 1) | | c&z | | c | | | | | | | z | |
| (1, 0) | | | | | | | | | | | | |
| (0, 2) | | | | | | | | | | | | |
| (0, 1) | | | | | | | | | | z | | |
| (0, 0) | | | | | z | | | | | z | | |

T P S P T V A H E S N W A K

Table interpretation
c or z means only c or zion in the table cell is matched in the spectrum. c&z means both c and z ions are matched.

Dynamic programming

**c**

Dynamic Programming Table (BestPath)

| (H, N) | | | | | | | | | | | | | |
|---|---|---|---|---|---|---|---|---|---|---|---|---|---|
| (2, 2) | 0 | 0 | 2 | 3 | 5 | 7 | 9 | 11 | 13 | 15 | 17 | 18 | 18 |
| (2, 1) | 0 | 0 | 0 | 0 | 1 | 2 | 2 | 2 | 2 | × | × | × | × |
| (2, 0) | × | × | × | × | × | × | × | × | × | × | × | × | × |
| (1, 2) | × | × | × | × | × | × | × | × | × | × | × | × | × |
| (1, 1) | 0 | 2 | 2 | 3 | 3 | 3 | 3 | 3 | 3 | × | × | × | × |
| (1, 0) | × | × | × | × | × | × | × | × | × | × | × | × | × |
| (0, 2) | × | × | × | × | × | × | × | × | × | × | × | × | × |
| (0, 1) | 0 | 0 | 0 | 0 | 0 | 1 | 1 | 1 | 1 | × | × | × | × |
| (0, 0) | 0 | 0 | 0 | 0 | 1 | 0 | 0 | 0 | 0 | × | × | × | × |

T P S P T V A H E S N W A K

Table interpretation
BestPath score of each cell is calculated by the best cell score of its left column plus its c/z ion number in the ScoreTable.

In this cell:
BestPath score is 13 + 2 = 15, where 13 is the best score of its left column, and its matched c/z ion number in ScoreTable is 2

× refers to the invalid cell

Backtracking the best path(s)

**d**

Dynamic Programming Table (BestPath)

| (H, N) | | | | | | | | | | | | | |
|---|---|---|---|---|---|---|---|---|---|---|---|---|---|
| (2, 2) | 0 | 0 | 2 | 3 | 5 | 7 | 9 | 11 | 13 | 15 | 17 | 18 | 18 |
| (2, 1) | 0 | 0 | 0 | 0 | 1 | 2 | 2 | 2 | 2 | × | × | × | × |
| (2, 0) | × | × | × | × | × | × | × | × | × | × | × | × | × |
| (1, 2) | × | × | × | × | × | × | × | × | × | × | × | × | × |
| (1, 1) | 0 | 2 | 2 | 3 | 3 | 3 | 3 | 3 | 3 | × | × | × | × |
| (1, 0) | × | × | × | × | × | × | × | × | × | × | × | × | × |
| (0, 2) | × | × | × | × | × | × | × | × | × | × | × | × | × |
| (0, 1) | 0 | 0 | 0 | 0 | 0 | 1 | 1 | 1 | 1 | × | × | × | × |
| (0, 0) | 0 | 0 | 0 | 0 | 1 | 0 | 0 | 0 | 0 | × | × | × | × |

T P S P T V A H E S N W A K

Table interpretation
If there is only one best path, then the sites are uniquely localized.

If there are multiple paths reaching the same best score, sites from the branch position to the merging position are regarded as a "site-group". Purple lines in this table indicate a site-group from S3 to T5, with glycan "H(1)N(1)"

Supplementary Note Figure 4. Detailed interpretation of the pGlycoSite algorithm for Fig. 2d-f in the main manuscript.

**Supplementary Note 4. Methods for sample preparation and LC-MS/MS**

**4.1 Standard N-glycopeptide preparation (Fig. 2c)**

Three synthetic standard N-glycopeptides: NVN[H(5)N(4)]ISYTVNDSFFPQRPQK, NVNISYTVN[H(5)N(4)]DSFFPQRPQK, and NVN[H(5)N(4)]ISYTVN[H(5)N(4)]DSFFPQRPQK, were kindly provided by Prof. Wei Huang (Shanghai Institute of Materia Medica, Chinese Academy of Sciences) through their previously developed methods[16, 17]. The three synthetic glycopeptides were suspended in 0.1% FA, mixed with equal amount and subjected to MS/MS analysis using PRM.

**4.2 Standard protein preparation (Fig. 2d)**

The standard glycoproteins IGM (SigmaAldrich, St. Louis, MO, U.S.A.) were dissolved in 50 mM NH4HCO3. The proteins were then reduced with 10 mM DTT at 37°C for 1 h and alkylated with 20 mM IAA for 0.5 h at room temperature in the dark. Then the proteins were divided into two aliquots. One is digested with trypsin and chymotrypsin. The other is digested with trypsin and Glu-C according to user manual. Then the two aliquots were desalted and lyophilized, respectively, and stored at −80 °C until subsequent glycopeptide enrichment.

Glycopeptides were enriched using a homemade zwitterionic hydrophilic interaction micro-column. Briefly, the desalted peptides of 1 mg were resuspended in 300 µL loading buffer containing 80% ACN and 1% TFA and then loaded onto a homemade micro-column containing 50 mg of ZIC-HILIC particles (Merck Millipore, Darmstadt, Germany) packed onto a C8 disk. The flow through was collected and reloaded onto the column for additional four times. Then, the column was washed with 200 µL loading buffer for four times, and finally eluted with 140 µL 0.1% TFA. The elution was collected and dried by vacuum centrifugation.

**4.3 inhibitor-initiated homogenous mucin-type O-glycosylation (IHMO) cell line preparation (Fig. 3a~3c)**

In IHMO approach, a kind of O-glycan elongation inhibitor, benzyl-N-acetyl-galactosaminide (GalNAc-O-bn), was applied to truncate the O-glycan elongation pathway during cell culture, generating cells with only truncated HexNAc(1) or HexNAc(1)NeuAc(1) O-glycans. It allows straightforward isolation of HexNAc-glycopeptides from cell lysates using lectin enrichment and acquisition of raw data through liquid chromatography-mass spectrometry (LC-MS/MS) with high energy collision dissociation (HCD) and electron-transfer/high energy collision dissociation (EThcD) fragmentation.

**Reagents**

Vicia Villosa Lectin (VVL) was purchased from Vector laboratories (SFO, USA). The peptide N-glycosidase F was obtained from NewEngland Biolabs (MA, U.S.A.). Antibodies for T, Tn and STn and all materials for cell culture were bought from Thermo Fisher Scientific (Waltham, MA, USA). Cell lines including HEK-293, Jurkat Clone E6-1, Hep 3B, MCF, and HeLa were obtained from National Collection of Authenticated Cell Cultures (Shanghai, China).

**Cell culture and inhibition O-GalNAcyaltion extending**

Cells were cultured according to related culture conditions (https://www.cellbank.org.cn/). When the cells grew to 70%, half of the cell dishes were added with 10 mM benzyl-N-acetyl-galactosaminide (GalNAc-O-bn) and cultured to another 48 hours. The treated HEK-293 cells and controls were labeled by T, Tn and STn antibodies, respectively according to antibody operation manual and then subjected to laser confocal microscopy, the image was shown in Supplementary Note Figure 5.

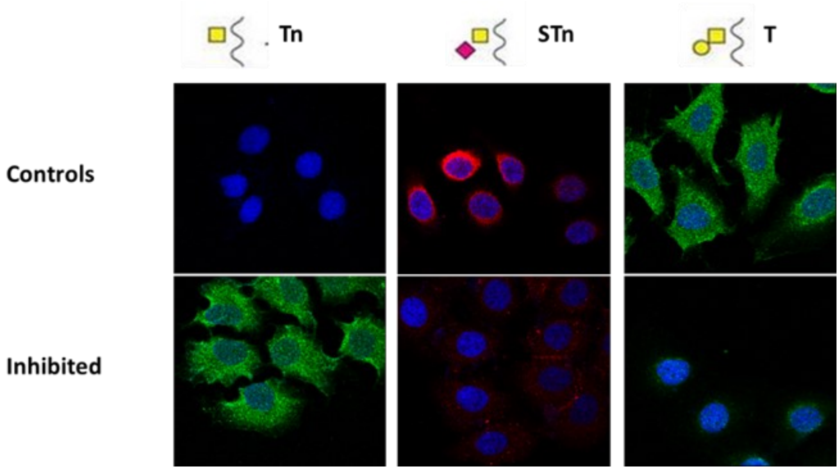

Supplementary Note Figure 5. Laser confocal microscopy images of controls and the inhibited O-glycopeptides using IHMO.

**Cell collection and Protein extraction**

The cells were collected in a tube after wash three times with PBS. Then the collected cells were lysed in fivefold-volume lysis buffer (4% SDS, 0.1M Tris/HCl, pH 8.0) with protease inhibitor (1mM PMSF, 1mM cocktail), followed by boiling at 100 °C for 10 minutes, ultrasonication for 20 minutes and centrifugation at 12, 000 g at 18 °C for 30 minutes to collect protein extracts. The protein concentration was determined by BCA method.

**Protein digestion**

Proteins were then reduced in 10mM dithiothreitol at 57 °C for 30 minutes, and then alkylated in dark by 20mM iodoacetamide at room temperature for 30 minutes. After carbamidomethylation, six volumes of acetone were added to precipitate the proteins at −20 °C for at least 3 hours. The precipitates were dissolved in a denaturing buffer (8M urea in 50mM NH4HCO3) following a ten-fold dilution with 50mM NH4HCO3. Trypsin was added to a final enzyme-to-substrate ratio of 1:50 and incubated at 37 °C overnight. The reactions were terminated by adding 0.5% trifluoroacetic acid. The peptide N-glycosidase F was added to the tryptic digested peptides with the ratio of 500 U/1 mg proteins, and incubated at 37 °C overnight to remove most of the N-glycans. Finally, all digested samples were centrifuged at 16,000 × g for 10 min and the supernatants were desalted using C18 column (Waters). The desalted peptides were then dried by vacuum centrifugation and used for glycopeptide enrichment.

**Glycopeptide enrichment**

The truncated O-glycopeptides were enriched with VVL lectin by the previously developed N-glyco-FASP method[18]. Briefly, about 1mg of lyophilized digests were resuspended in 400 µL binding buffer (40 mM Tris-HCl pH 7.6, 2mM MnCl2, 2mM CaCl2, 1mM NaCl), transferred into a filter unit (30, 000MWCO). Then, 500 µg VVL was added to the filter unit containing peptides, mixed at 550 rpm in a thermo-mixer for 1 minute and incubated without mixing for 60 minutes at room temperature. After that, centrifuge the filter unit at 14,000 x g for 10 minutes, add 200 µL binding buffer and centrifuge the filter unit again to wash the non-specific binding peptides with two replicates. Then, transfer the filter unit to a new collection tube, add 400 µL 200 mM N-acetylgalactosamine in binding buffer and incubate at room temperature for 45 minutes. Finally, the eluting solution containing enriched glycopeptides were collected, desalted with C18 Sep-Pak and lyophilized for further LC-MS/MS analysis.

**4.4 Human serum preparation (Fig. 3e~3f)**

Informed consent was obtained under protocols that were approved by an institutional review board. The research followed the tenet of the Declaration of Helsinki and was approved by the Ethics Committee of the Fudan University. A total of ten healthy human blood samples were collected, immediately placed on ice for 30 minutes and then centrifuged at 2000 g for 15 minutes. The supernatant was collected. A pooled serum specimen mixed from equal volume of the supernatant was used in this study. A volume of 15 µL of the pooled serum was added to 185 µL of 50 mM NH4HCO3, and placed in boiling water and ice 40 times alternately to denature the proteins. The proteins were then reduced with 10 mM DTT at 37°C for 1 h and alkylated with 20 mM IAA for 0.5 h at room temperature in the dark. Then the proteins were digested into peptides and desalted using Sep-Pak C18 cartridges (Waters, USA). The desalted peptides were then dried by vacuum centrifugation. For the detection of O-glycosylation, the peptide N-glycosidase F (PNGase F, glycerol free, 500 U μL-1), which was obtained from NewEngland Biolabs (MA, U.S.A.), was added to the tryptic digested peptides with the ratio of 500 U/1 mg proteins, and incubated at 37 °C overnight to remove most of the N-glycans. Then the peptides were desalted, lyophilized, and stored at −80 °C until subsequent glycopeptide enrichment.

**Glycopeptide enrichment**

Glycopeptides were enriched using a homemade zwitterionic hydrophilic interaction micro-column. Briefly, the desalted peptides of 1 mg were resuspended in 300 μL loading buffer containing 80% ACN and 1% TFA and then loaded onto a homemade micro-column containing 50 mg of ZIC-HILIC particles (Merck Millipore, Darmstadt, Germany) packed onto a C8 disk. The flow through was collected and reloaded onto the column for additional four times. Then, the column was washed with 200 μL loading buffer for four times, and finally eluted with 140 μL 0.1% TFA. The elution was collected and dried by vacuum centrifugation.

**4.5 Preparation of fission yeast and budding yeast for aH-glycopeptide analysis (Fig. 4c)**

**Sample preparation**

Washed fission yeast and budding yeast were snap-frozen in liquid nitrogen, ground to a fine powder with mortar and pestle in liquid nitrogen, and stored at -80°C until use. The grinding powder from different organisms were processed to protein extraction and digestion. The powder was dissolved in fivefold-volume lysis buffer (4% SDS, 0.1M Tris/HCl, pH 8.0) with protease inhibitor (1mM PMSF, 1mM cocktail), followed by boiling at 100 °C for 10 minutes, ultra-sonication for 5 minutes and centrifugation at 12, 000 g at 18 °C for 30 minutes to collect protein extracts. The protein concentration was determined by BCA method.

**Protein digestion**

Proteins were then reduced in 10mM dithiothreitol at 57 °C for 30 minutes, and then alkylated in dark by 20mM iodoacetamide at room temperature for 30 minutes. After carbamidomethylation, six volumes of acetone were added to precipitate the proteins at −20 °C for at least 3 hours. The precipitates were dissolved in a denaturing buffer (8M urea in 50mM $NH_4HCO_3$) following a ten-fold dilution with 50mM $NH_4HCO_3$. Trypsin was added to a final enzyme-to-substrate ratio of 1:50 and incubated at 37 °C overnight. The reactions were terminated by adding 0.5% trifluoroacetic acid. Finally, all digested samples were centrifuged at 16,000 × g for 10 min and the supernatants were desalted using C18 column (Waters). The desalted peptides were then dried by vacuum centrifugation and used for glycopeptide enrichment.

**Glycopeptide enrichment**

Glycopeptides were enriched by zwitterioic hydrophilic interaction liquid chromatography (ZIC-HILIC) method. Briefly, the desalted peptides of 1 mg were resuspended in 300 µL loading buffer containing 80% acetonitrile and 1% trifluoroacetic acid and then loaded onto an in-house micro-column containing 50 mg of ZIC-HILIC particles (Merck Millipore) packed onto a C8 disk. The flow through was collected and reloaded onto the column for additional four times. Then, the column was washed with 200 µL loading buffer for four times. Finally, the glycopeptides that have been enriched in the column were collected by eluting with 600 µL 0.1% trifluoroacetic acid and dried by vacuum centrifugation.

**4.6 Preparation of N-glycome analysis on fission yeast (Supplementary Fig. 3)**

We set up a MS-based glycomic analytical pipeline to identify the N-glycome of fission yeast to verify the existence of "Hex+17" in yeast samples.

**Chemicals and reagents**

Sequencing grade trypsin was purchased from Hua LiShi Scientific Corporation (Beijing, China). PNGase F (glycerol free) was purchased from New England Biolabs (Ipswich, MA). The BCA protein assay kit was purchased from Pierce (Rockford, IL). SEP-PAK C18 columns were obtained from Waters (MA, USA). Porous graphitized carbon (PGC) columns were purchased from Grace (Columbia, MD). Ammonium bicarbonate was purchased from Sinopharm Chemical Reagent (Shanghai, China). All other reagents were obtained from Sigma-Aldrich (St. Louis, MO).

**Purification of N-glycans**

Fission yeast were snap-frozen in liquid nitrogen, ground to a fine powder with mortar and pestle in liquid nitrogen, and stored at -80°C until use. The grinding powder from yeast were processed to protein extraction and digestion. The powder was dissolved in fivefold-volume lysis buffer (4% SDS, 0.1M Tris/HCl, phoH 8.0) with protease inhibitor (1mM PMSF, 1mM cocktail), followed by boiling at 100 °C for 10 minutes, ultrasonication for 5 minutes and centrifugation at 12, 000 g at 18 °C for 30 minutes to collect protein extracts. The protein concentration was determined by BCA method.

Proteins were reduced in 10 mM dithiothreitol at 37 °C for 60 minutes, and then alkylated in dark by 20 mM iodoacetamide at room temperature for 30 minutes. After carbamidomethylation, 6 volumes of acetone were added to precipitate the proteins at -20 °C for at least 3 hours. The precipitates were dissolved in 50 mM NH4HCO3. Trypsin was added to a final enzyme-to-substrate ratio of 1:50 and incubated overnight at 37 °C. The reactions were terminated by heating at 100 ℃ for 10 minutes. Then, the tryptic digestion samples were added with PNGase F (20 U/100 μg glycoproteins) and incubated at 37 °C overnight for glycan release. Finally, the mixtures were centrifuged at 14,000 x g for 10 min. The supernatants were purified using SEP-PAK C18 column. The wash flows were collected and desalted by PGC columns. Purified glycans were lyophilized through vacuum centrifugation and dissolved in 0.1% formic acid for LC-MS/MS analysis.

**Targeted N-glycan search**

After N-glycopeptide MS/MS data were identified by pGlyco3, all identified unique N-glycan compositions including aH-glycans were compiled into an N-glycan list. The N-glycan spectra were searched against the N-glycan list using the targeted approach and scored by the glycan B/Y ion-counting score. Only the top-scored glycan was kept for each spectrum, and the results were further filtered by score ≥ 10. The identified N-glycopeptides as well as the aH-glycopeptides were validated by the filtered N-glycan search results.

#### 4.7 EThcD-MS/MS for O-Man glycopeptide analysis on fission yeast (Supplementary Fig. 4)

We conducted experiments for intact O-glycopeptide identification in fission yeast. The whole experimental procedures of sample preparation are the same with that for glycomic analysis on fission yeast. O-glycopeptides were enriched using ZIC-HILIC with the same condition for N-glycopeptide enrichment from the tryptic digests, expect for the treatment with PNGase F (20 U/100 μg glycoproteins) to release N-glycans before enrichment.

#### 4.8 LC-MS/MS analysis for samples prepared in 4.1 to 4.7

All LC-MS/MS analyses were performed on a nanospray LC-MS/MS on an Orbitrap Fusion Tribrid system (Thermo Fisher Scientific, Waltham, MA, U.S.A.) equipped with an EASY-nLC TM1100 system (Thermo Fisher Scientific, Waltham, MA, USA). Solvent A was a 0.1% FA aqueous solution. Solvent B was ACN containing 0.1% FA. Detailed LC-MS/MS parameters are listed in Supplementary Data 2.

### Supplementary Note 5. Comprehensive comparisons and analyses of glycopeptide identifications based on PXD005565

#### The rationality of yeast glycopeptides as benchmarks

Deep investigation on the differences between different search engines such as pGlyco3 and Byonic is very useful to indicate the future improvement for each tool. However, this kind of investigation is difficult as they have different MS1 precursor detection, peptide scoring, glycan scoring, and FDR filtering strategies. And more importantly, there are too few large-scale benchmark glycopeptide datasets in the glycoproteomics field. And without "ground truth", the comparisons can hardly point out which direction we could make efforts to improve our software tools. Fortunately, yeast only contains high-mannose N-glycans (glycans with composition H(X)N(2)), making it become a good glycan-level benchmark. Maybe one concern is that the complexity of high-mannose N-glycans is too low for mammalian systems. But the presence of aH in yeast samples makes the complexity much higher, as shown in Supplementary Note Figure 6. The complexity of human N-glycans is not as high as we expected because most of the Y-complementary compositions could be uniquely determined by their masses within ±0.03 Da. For ±0.04 Da tolerance, the uncertainty is introduced because "H(7)=1134.370 Da" and "N(2)A(2)F(1)=1134.407 Da", and it will be lower if we check the NeuAc diagnostic ions. aH-glycans are more difficult to be distinguished from non-aH-glycans if search engines only score against small Y ions such as core Y-ions as their Y-complementary ions may be too large to be uniquely determined. The main uncertainty is owning to "H(3)aH(2)=844.317 Da" and "N(2)F(3)=844.332 Da", hence "H(X)N(Y)aH(2)" are easily identified as "H(X-3)N(Y+2)F(3)". Ambiguous Y-complementary ions might easily result in inaccurate glycan identifications. Therefore, from the glycan identification point of view, the yeast sample with unexpected aH is complex enough as one of the glycan-level benchmarks. Therefore, all the following analyses were based on the fission yeast dataset (PXD005565).

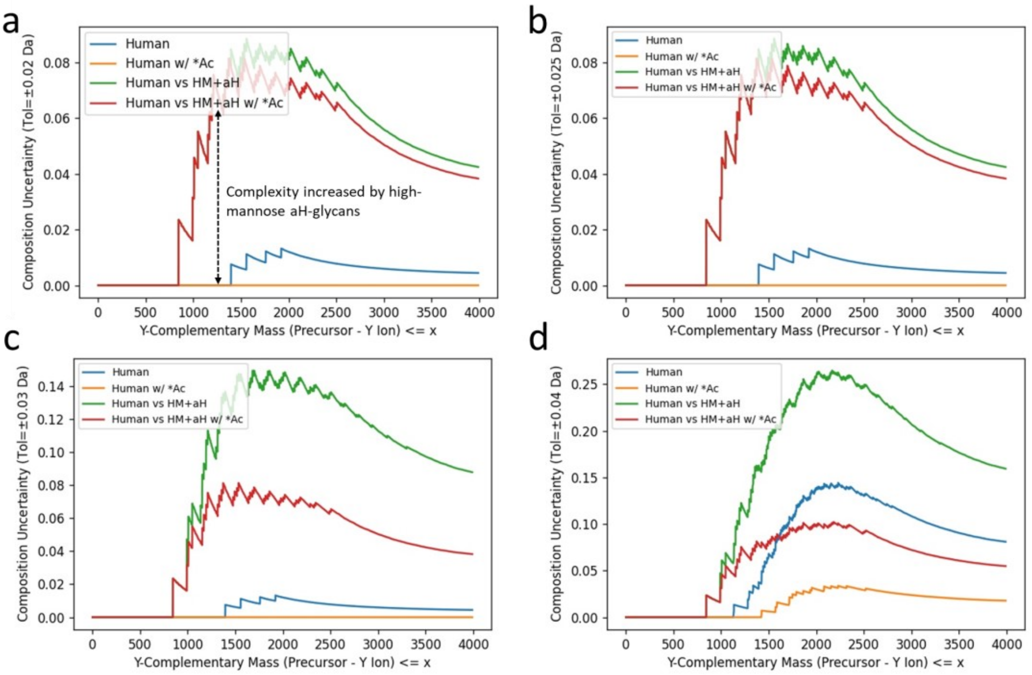

Supplementary Note Figure 6. High-mannose (HM) aH-glycopeptides increase the complexity of yeast N-glycans. For a given glycan database, all theoretical Y-complementary ions are calculated. If a Y-complementary mass could not uniquely determine a glyco-composition within a given mass tolerance (a: ±0.02 Da, b: ±0.025 Da, c: ±0.03 Da, d: ±0.04 Da), we say it is uncertain. The uncertainty is defined as the proportion of uncertain Y-complementary masses for all Y-complementary masses not larger than x. The uncertainty reflects the complexity of a glycan database. "Human" refers to the Y-complementary ions from the human N-glycan database; "w/ *Ac" means using NeuAc-diagnostic ions to distinguish NeuAc-glycans; "Human vs HM+aH" refers to human N-glycan database mixed with high-mannose aH-glycans. All curves go down after 2500 Da because there are no many large (>2500 Da) glycans in the database, hence they seem easier to be uniquely identified. But in fact, large masses would share more compositions.

This does not mean that human glycans are easy to identify even if human glycans could be uniquely determine by accurate masses, a key issue is that aH-glycopeptides are quite commonly observed in different datasets (Supplementary Figure 8). And unexpected factors would influence the identifications as well, some of them will be discussed below, and some others had been well discussed previously[19, 20].

### How different glycan databases influence the glycan identification?

The fission yeast data (PXD005565) were searched by pGlyco3 with different glycan databases (182 GDB, 1234 GDB and 1234 GDB+aH), and the results showed very good consistencies (Supplementary Note Figure 7a). Since pGlyco3 only searched glycan structures, we extracted all glycan structures in the pGlyco-N-Human glycan database if their compositions were found in the 182-N-glycan database. There were only 10 compositions in 182-N-Glycan we not found in pGlyco-

N-Human. On PXD005565 yeast dataset, with 182-N-Glycan database, pGlyco3 could obtain 3255 GPSMs, of which 26 GPSMs were not identified as high-mannose glycans, indicating 0.8% (26/3255) glycan-level FDR, as shown in Supplementary Note Figure 7b. When 1234 GDB was used, the non-high-mannose ratio of the GPSMs was 4.0% (135/3405, Supplementary Note Figure 7b). 111 of 135 false identifications were because the "H(X)N(2)aH(2)"-glycans were incorrectly identified as "H(X-3)N(4)F(3)". 182 GDB did not have such an issue as there were no "H(X-3)N(4)F(3)"-glycans, showing that inappropriately increasing the glycan database might potentially increase the identification FDR. But considering aH modification for 1234 GDB (pGlyco-N-Mouse) could decrease the number of "H(X-3)N(4)F(3)"-glycans from 111 to 5, indicating the importance of aH-search if aH was presented in the sample. But aH search had its own problems. There were 1.9% (114/5893, 1.9% glycan-level FDR) non-high-mannose GPSMs when searching with 1234 GDB+aH. We found that 72 of 114 false identifications were because of "F(1)aH(1) – H(2) = 1 Da". That means pParse (the MS1 precursor detection module of pGlyco3) incorrectly exported M1 as M0 for these spectra, an MS1 XIC example was shown in Supplementary Note Figure 10.

These analyses showed that: (1) glycan database is not the larger the better, inappropriately increasing the database would result in more false identifications; (2) if aH is presented in the sample, it is better to add aH into the search space, otherwise it may lead to incorrect identifications. Fortunately, with the glycan-first search and glycan quality control, pGlyco3 does not appreciably increase the glycan error rates with large glycan databases.

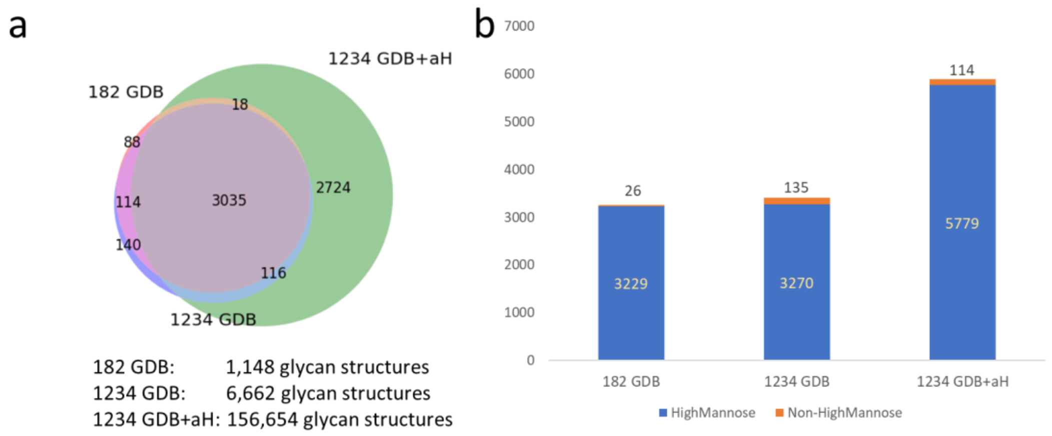

Supplementary Note Figure 7. Searching PXD005565 using different glycan databases with pGlyco3. (a) Venn diagram of three search results; (b) Numbers of non-high-mannose GPSMs. 182 GDB: N-glycan structure database designed from 182-N-Glycan database of Byonic; 1234 GDB: pGlyco-N-Mouse glycan database; 1234 GDB + aH: pGlyco-N-Mouse glycan database with aH.

### Comparisons of Byonic and pGlyco3 based on PXD005565

Taking the high-mannose glycans as the "ground truth" in the yeast samples, we investigated why pGlyco3 and Byonic obtained different results on the same MS2 spectra. For better investigation, we use the results of "1234 GDB+aH" in these comparisons to discover different situations. Firstly, we classified the different results into 4 classes, as show in Supplementary Note Figure 8. 1. "Different Peptides": peptides were different; 2. "pGlyco3 True": glycans were different but pGlyco3 identified high-mannose glycans; 3. "Byonic True": glycans were different but Byonic identified high-mannose glycans; 4. "Both False": both identified false glycans. There were 120 MS2 scans inconsistently identified, including 9 "Different Peptides", 77 "pGlyco True", 33 "Byonic True", and 1 "Both False" results. Peptide-level identification of Byonic and pGlyco3 were very consistent as there were only 9 (9/2246 = 0.4%) different peptides, indicating high peptide-level accuracies for both Byonic and pGlyco3. We then manually checked "pGlyco3 True" identifications, and found out that 88% (68/77) of Byonic's N-glycans contained NeuAc. These false identifications were obtained because of "H(5)aH(1) – N(2)A(2) = 1 Da" and "H(3)aH(2) – N(2)A(1)F(1) = 1 Da". But these errors could be removed if we further checked the existence of NeuAc by its diagnostic ions and hence increase the accuracy for Byonic. In fact, among all 323 (323/3406 = 9.5% of all GPSMs) non-high-mannose identifications of Byonic, 73.7% (238/323) incorrect identifications could be removed by checking NeuAc-diagnostic ions, resulting in 2.9% glycan-level FDR which is quite acceptable, showing the importance of the glycan-level quality control for Byonic. We further analyzed MS1 information for Byonic identifications, as shown in Supplementary Note Figure 9, no M0 (monoisotopic peak) signals of Byonic's result were found in adjacent retention time of the searched MS2 scan. After we manually checked several MS1 XICs of these 68 spectra with NeuAc-glycans, we speculated that there might be some issues in the MS1 precursor detection (peak picking) module of Byonic. For 33 "Byonic True" identifications in pGlyco3, 27 of them were due to "F(1)aH(1) – H(2) = 1 Da" as discussed previously, and an MS1 example were shown in Supplementary Note Figure 10. For uniquely identified spectra of pGlyco3, the glycan-level FDR would be 2.2% (80/3647). The main errors of these 80 GPSMs were because "H(X)N(2)aH(2)" was incorrectly identified as "H(X-3)N(4)F(3)". Byonic had no "H(X-3)N(4)F(3)"-identifications because there are no "H(X-3)N(4)F(3)"-glycans in the 182-N-glycan database. But it was a potential problem for all software tools if a larger glycan database (e.g., 1234 GDB) was used.

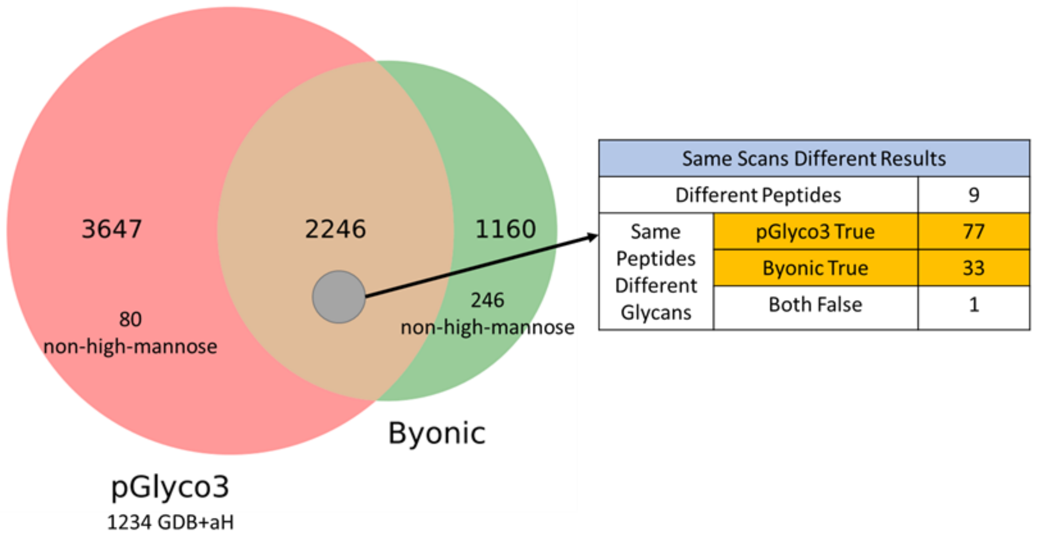

Supplementary Note Figure 8. Different results of pGlyco3 (1234 GDB+aH) and Byonic on the same MS2 spectra. “non-high-mannose” means the number of spectra that were not identified as high-mannose glycans.

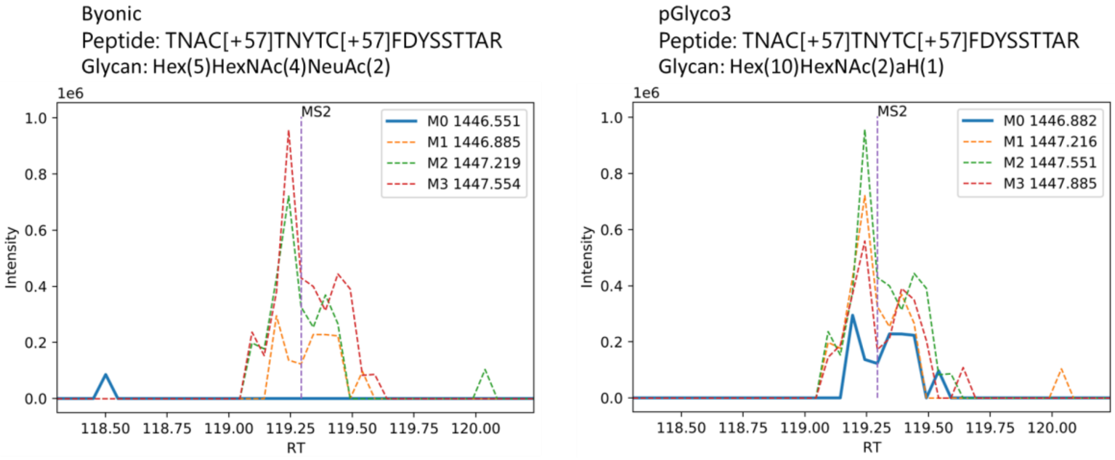

Supplementary Note Figure 9. The MS1 XIC of a “pGlyco True” example for Byonic and pGlyco3 identifications. M0 is the monoisotopic m/z.

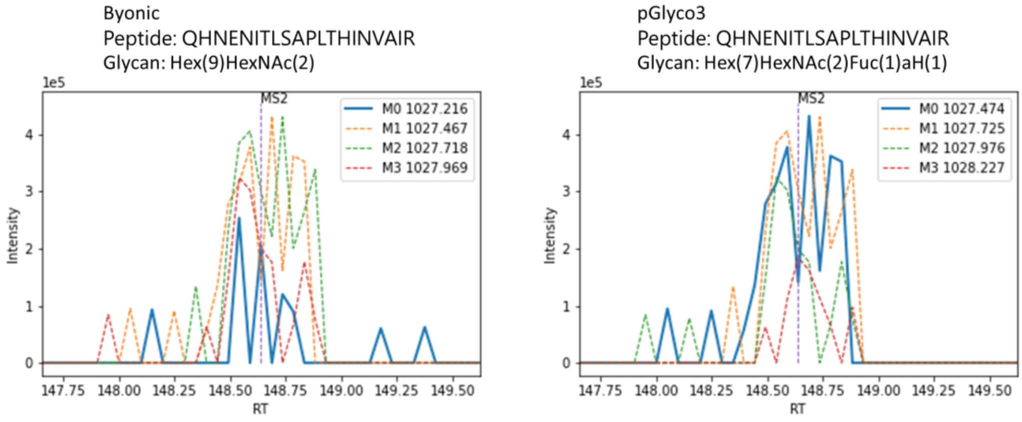

Supplementary Note Figure 10. The MS1 XIC of a “Byonic True” example for Byonic and pGlyco3 identifications. M0 is the monoisotopic m/z. pParse sometimes fails to handle unstable MS1 signals, which could be well processed by

Byonic.

Other than diverse precursor detection and search space problems, different glycan and peptide scoring and FDR filtration strategies would also result in different FDR ranks for same GPSMs, those are also the reasons why both Byonic and pGlyco3 lost some correct identifications. For example, the GPSM displayed in Supplementary Note Figure 11 was identified by Byonic but lost by pGlyco3 because it was at 1.5% peptide FDR in pGlyco3; and the GPSM displayed in Supplementary Note Figure 12 was identified by Byonic but lost by pGlyco3 because it was at 10% glycan FDR in pGlyco3. The recent community study also showed that, even for Byonic itself, the identifications were quite diverse searched by different teams[21].

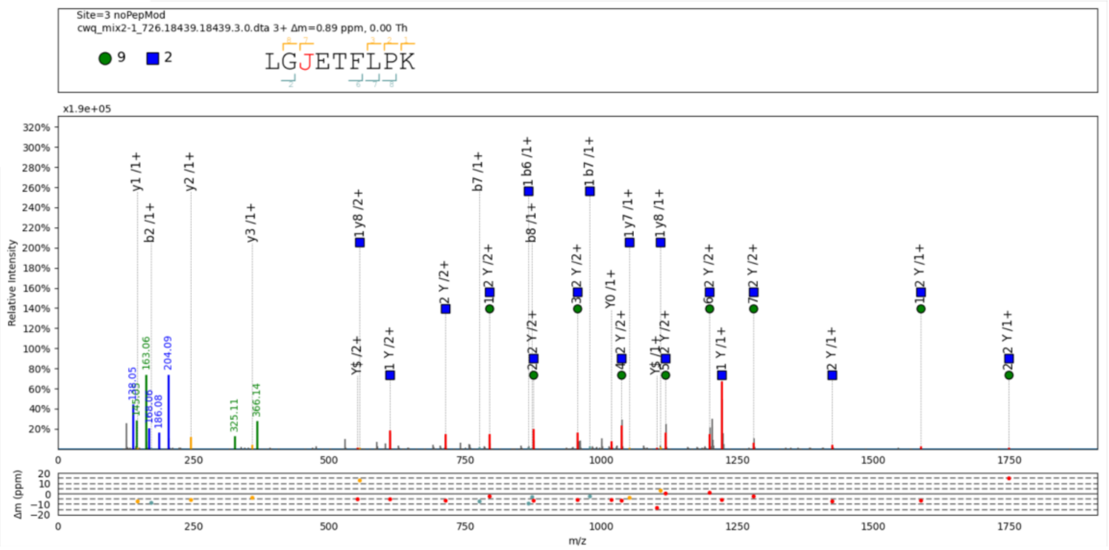

Supplementary Note Figure 11. A GPSM identified by Byonic but lost by pGlyco3 because it is at 1.5% peptide FDR in pGlyco3. The glycan annotation is based on pGlyco3's Y ions.

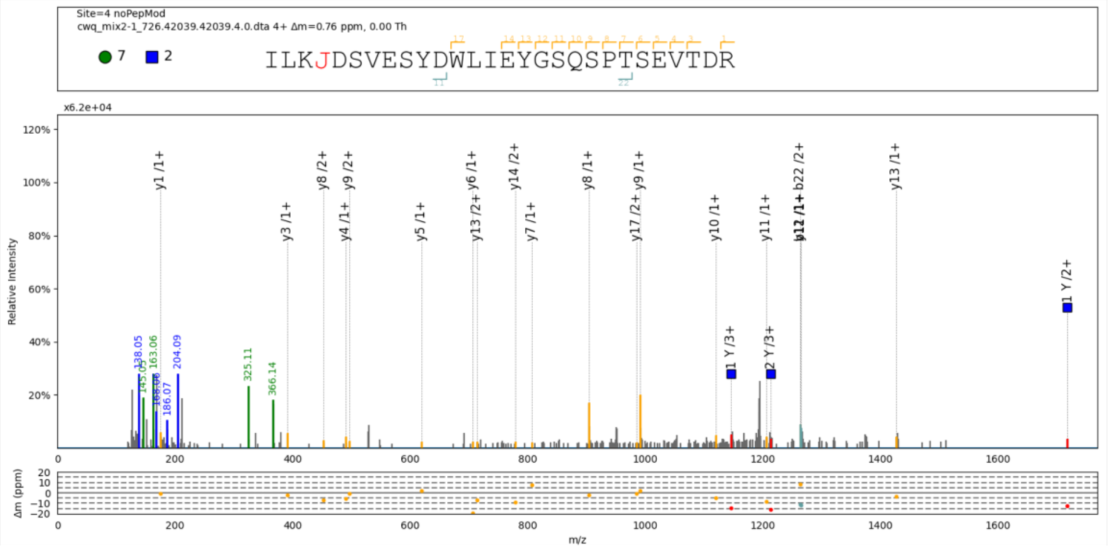

Supplementary Note Figure 12. A GPSM identified by Byonic but lost by pGlyco3 because it is at 10% glycan FDR in pGlyco3. The glycan annotation is based on pGlyco3's Y ions.

As a conclusion, in the routine glycoproteomic workflow, as we do not know how many aH-glycopeptides are presented in the samples, the pros and cons should be kept in mind if we consider aH search or not. For both pGlyco3 and Byonic, and maybe other tools as well, these analyses suggest how to further increase the glycan identification confidence:

1. Increasing mass detection (instruments) and searching (software) accuracies of MS1/MS2 would be the most helpful;
2. More accurate precursor detection algorithm is needed;
3. Checking the glyco-diagnostic ions could decrease the uncertainty of glycan assignments. It is very useful to control the glycan FDR for Byonic, hence users can consider it in the post-processing step or downstream analysis;
4. Producing Y ions as many as possible to cover large Y ions, and hence reduce the biggest Y-complementary masses and increase the confidence of glycan assignments, this is why sceHCD is recommended (lower CE for large Y ions);
5. Considering as large Y ions as possible in the scoring functions. Penalize the glycan if many of its Y ions could not be matched;
6. Sample-specific glycan databases would be helpful, though it is difficult to design such a glycan database for each sample from mammalian systems. With expert knowledge, users are able to design their own glycan databases with and without modified saccharide units for pGlyco3 using GlycoWorkbench. Software tools also need to provide different built-in glycan databases for users to choose and test, that is what we are going to do in the next step by working with glycobiology experts.
